# Latent defense response to non-pathogenic microbial factors impairs plant-rhizobacteria mutualism

**DOI:** 10.1101/2020.04.24.058990

**Authors:** Yu Yang, Shenglan Chen, Li Peng, Xiaomin Liu, Richa Kaushal, Fengtong Yuan, Sunil K. Singh, Danxia He, Suhui Lv, Juan I. Vílchez, Rafael J. L. Morcillo, Wei Wang, Weichang Huang, Mingguang Lei, Chun-Peng Song, Jian-Kang Zhu, Paul W. Paré, Huiming Zhang

## Abstract

Unlike pathogens that trigger plant defense responses, commensal or beneficial microbes are compatible with plants and do not elicit a defense response. An assumption underlying the compatibility is that plants are inert in mounting a defense response to non-pathogenic microbial factors. However, the mechanisms underlying this inertness in defense are unknown. Here a forward genetic screen led to the isolation of an Arabidopsis mutant displaying a new type of immunity which we named as latent defense response (LDR) to a beneficial rhizobacterium. The mutant, known as *gp1* for *Growth-Promotion 1*, is impaired in rhizobacteria-induced plant growth-promotion due to disrupted oleic acid homeostasis and consequent activation of defense responses. Several bacterial volatile compounds trigger LDR in *gp1* but not wild type plants. GP1 dysfunction strongly represses colonization of the beneficial rhizobacterium and alters root-associated microbiota. Our findings reveal a hidden layer of plant defense, LDR, which is suppressed by GP1 to allow mutualistic association between plants and beneficial rhizobacteria.

**One Sentence Summary:** A hidden layer of host immunity against non-pathogenic microbes leads to plant incompatibility with beneficial rhizobacteria.

## Introduction

Plants naturally live with a wide variety of soil microbes that can be pathogenic, commensal, or beneficial (*1–3*). While beneficial microbes can promote plant growth or improve plant tolerance to stresses, pathogens impose threats that in some cases can be deadly to plants (*4, 5*). Hence the detection of and responses to microbial factors are critical for plant survival. Microbes produce and secret a complex array of metabolites that all potentially can be perceived by neighboring plants (*6, 7*). Some microbial factors are perceived by plants as pathogenic and thereby trigger plant defense responses; meanwhile many others appear to be inert in that they do not trigger plant defense (*8–10*). This seemingly inert relation can be easily accepted but is vague, although it underlies the commensality or mutualism between plants and non-pathogenic microbes.

Plant growth-promoting rhizobacteria (PGPR) are plant-beneficial soil microbes that live within the rhizopshere and induce plant vigor as demonstrated by increased plant growth and/or stress tolerance (*4, 11*). PGPR have been successfully applied to crop species to increase seedling emergence, plant weight, crop yield, and stress tolerance (*12, 13*). These beneficial soil microbes induce plant vigor through production of one or multiple bacteria-derived factors including phytohormones such as auxin and cytokinin, the enzyme ACC (1-aminocyclopropane-1-carboxylate) deaminase that reduces plant ethylene levels, and siderophores that facilitate root uptake of metal nutrients (*4, 11*). In addition to these non-volatile compounds, some PGPR strains also release microbial volatiles compounds capable of affecting plant physiology under normal and stressed conditions (*11, 14*).

*B. amyloliquefaciens* strain GB03 is an established beneficial microbe to plants both in soil and in artificial medium (*11, 15, 16*). Previous studies have reported that GB03-produced microbial volatiles (hereafter referred to as GMVs) cause multiple beneficial effects on plants, such as enhanced lateral root development and increased contents of photosynthetic apparatus, through modulation of phytohormone homeostasis and nutrient uptake (*17–19*), although it remains unclear how plant biological processes were integrated by the microbial factors to produce the multiple beneficial traits. In addition, a recent study demonstrated that diacetyl, a GMV component, has opposing effects on GB03-plant association under phosphate-sufficient and phosphate-deficient conditions (*20*). These findings points to the importance to identify plant molecular factors that are indispensable for proper responses to PGPR.

## Results

### Isolation of an Arabidopsis mutant that shows defective inducible vigor

To identify molecular factors underlying plant inducible vigor, we searched for Arabidopsis mutants defective in showing growth-promotion triggered by *B. amyloliquefaciens* GB03. The forward genetic screen was performed with GMV-treated plants grown in petri dishes with 1/2-strength MS solid medium. The petri dishes contained partitions that physically separate the medium for plant growth from the medium for bacteria growth (Figure 1A), so that the bacteria can affect the plants only through airborne diffusion of the volatile emissions. We isolated a mutant that showed reduced growth promotion in response to GMVs, and named it *gp1-1* (*growth promotion 1-1*). At 11 days after treatments (DAT), GMVs induced 5.7 and 2.8 folds increases in the fresh weight of wild type (wt) plants and of the *gp1-1* mutant, respectively (Figure 1B). Quantification of plant leaf area showed a similar pattern that GMV-induced plant vigor is compromised in the *gp1-1* mutant (Figure S1A).

**Figure 1.**
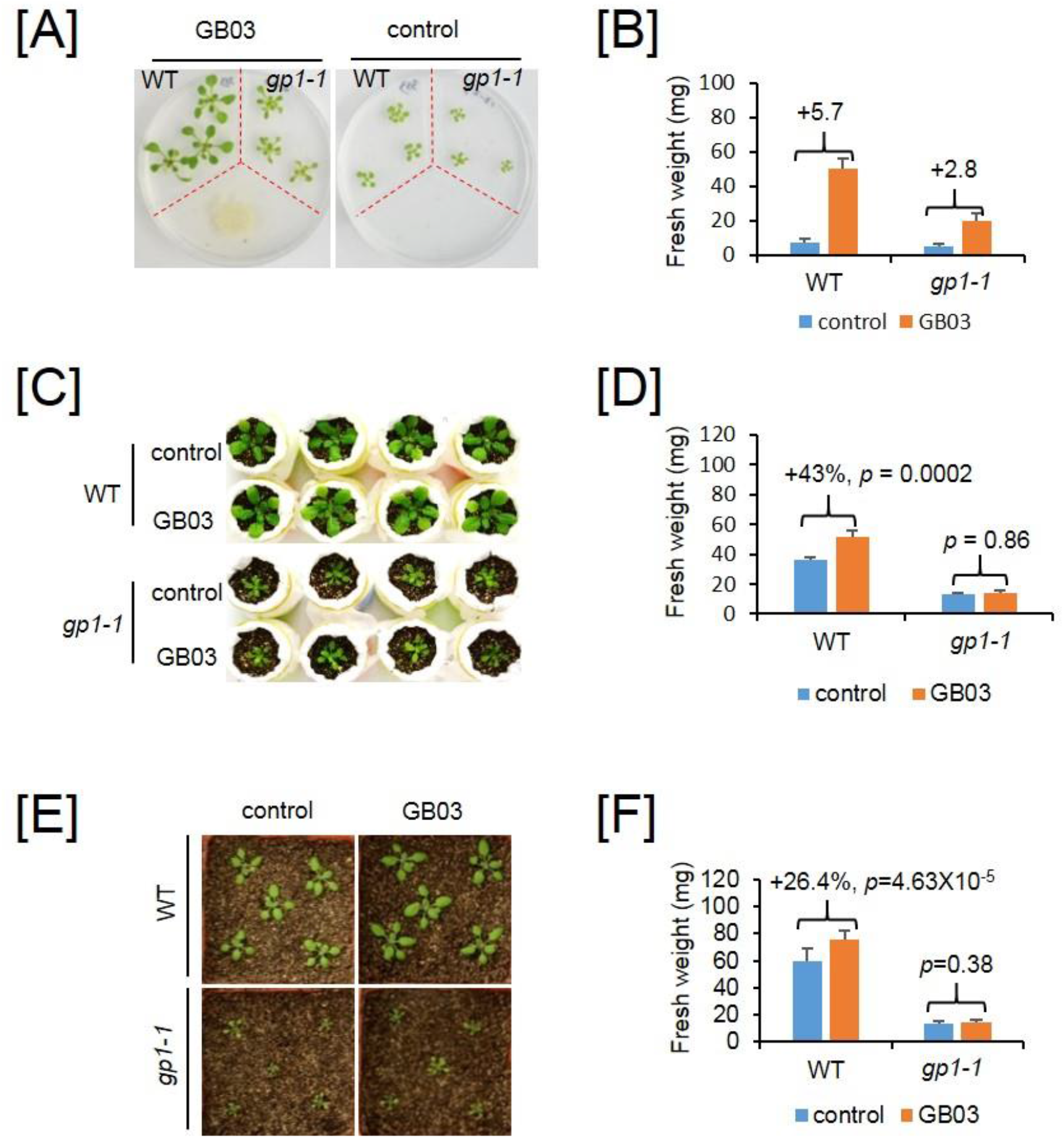
*gp1* mutation impairs GB03-induced growth promotion in Arabidopsis. The *gp1-1* mutant was compared with its wild type (WT) plants in different growth systems. **[A]** Plants were grown in 1/2-strength Murashige & Skoog medium with or without exposure to GB03-emitted microbial volatiles (GMVs). The petri dishes contained plastic partitions (red dotted lines), which separate the medium for plant growth and the medium for bacteria, so that the bacteria can affect the plant only through volatile emissions. Images were taken at 11 days after treatment (DAT). **[B]** Quantification of whole-seedling fresh weight of plants grown in 1/2-strength MS medium at 11 DAT. Values are mean ± SD (n = 6). **[C]** Plants were grown in a soil-in-tube system (see Figure S1B for elucidation) that allows GMVs to diffuse from the bottom of the 50mL tubes. Images were taken at 14 DAT. **[D]** Quantification of aerial-portion fresh weight of plants grown in the soil-in-tube system at 14 DAT. Mean ± SD (n = 5). **[E]** Plants were grown in soil with or without GB03 innoculation. Images were taken at 9 DAT. **[F]** Quantification of aerial-portion fresh weight of soil-grown plants with or without GB03 inoculation at 11 DAT. Mean ± SD (n = 12). Differences between the treated and the control plants were shown as fold changes above each group for all quantification of fresh weight. Statistical analysis are based on student t-test.

In addition to using the airtight petri dish system that appears to favor perception of GMVs by leaves, we also examined *gp1-1* responsiveness to GMVs in an open system, in which plants were grown in soil suspended above the bacteria (Figure S1B). In such an experimental setup, GMVs would diffuse through rhizosphere soil pores and be more likely perceived by roots. Exposure to GMVs resulted in 43% increase in the fresh weight of wt plants; in contrast, GMVs failed to promote growth of *gp1-1* (Figure 1C and D). Therefore, the *gp1-1* mutant is impaired in plant responsiveness to GMVs.

We further compared *gp1-1* with the wt plants in soil with direct bacteria inoculation to roots (Figure 1E). GB03 inoculation resulted in 29.4% and 26.4% increases in wt plant leaf area and fresh weight, respectively; in contrast, *gp1-1* showed no growth-promotion effects in the presence of GB03 (Figure 1F; Figure S1C). Altogether these results demonstrate an essential role of GP1 in plant vigor induced by GB03.

### The loss of inducible vigor in gp1 plants is due to gene dysfunction of At2g43710

The *gp1-1* mutant was backcrossed to its wt plant, and the resulting F1 progeny exhibited GB03-induced vigor similarly as wt plants, indicating that the mutation is recessive. The self-pollinated F2 progenies segregated with a ~3:1 ratio for wild type versus mutant regarding to the reduced growth-promotion phenotype, suggesting that the *gp1-1* mutation is controlled by a single nuclear gene. Map-based cloning was then performed and the mutation was mapped to chr2 in a mapping interval between 17 970Kb and 18 145Kb. Subsequently, whole genome sequencing revealed a C-to-T point mutation in the second exon of the gene At2g43710. This point mutation was predicted to cause an amino acid change from Thr155 to Ile (Figure S2A).

To confirm that the mutation of At2g43710 is responsible for the loss of inducible vigor, we performed gene complementation in the *gp1-1* mutant. The genomic DNA of At2g43710 including its native promoter was cloned and tagged with 3XHA, 3XMYC, or 3XFLAG, and the constructs were transformed into the *gp1-1* mutants. Western blots confirmed the protein accumulation of each tagged transgenic protein in the *gp1-1* background (Figure 2A; Figure S2B and C). As expected, transgenic expression of At2g43710 in the *gp1-1* mutant completely restored GMV-induced plant vigor, as shown by plant leaf area and fresh weight measurements (Figure 2B-C; Figure S2D-G). Meanwhile, a mutation caused by a T-DNA insertion (*gp1-2*) in the second exon of At2g43710 mimics the *gp1-1* mutant in suppressing GMV-induced plant vigor (Figure S3). Together, these results demonstrate that the loss of inducible vigor in *gp1* plants is due to gene dysfunction of At2g43710.

**Figure 2.**
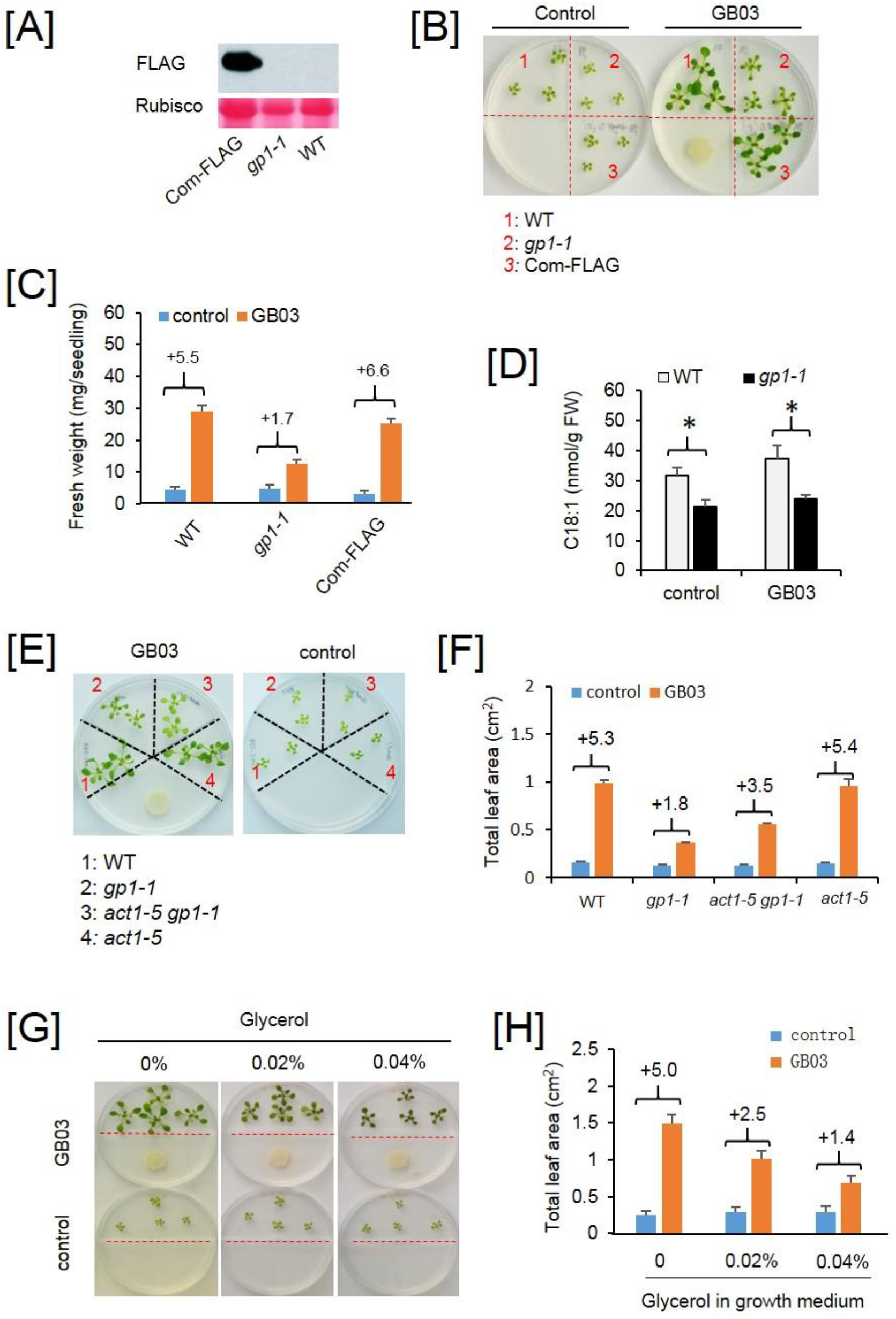
The impairment of inducible vigor in the *gp1* mutants is caused by At2g43710 dysfunction. [A] Transgenic expression of Pro_GP1_::GP1-FLAG in *gp1-1* (Com-FLAG for simplicity) restored protein accumulation of GP1. [B] Transgenic expression of Pro_GP1_::GP1-FLAG in *gp1-1* restored plant inducible vigor. Images were taken at 9 DAT. Red dotted lines indicate plastic partitions. [C] Whole-seedling fresh weight quantified at 9 DAT. Mean ±SD (n=6). Fold changes are shown above each group. [D] Oleic acid contents in WT and *gp1-1* with or without GB03 treatment. Samples were collected at 3 DAT. Mean ± SE (n=3). * p < 0.05, student t-test; [E] Comparison of plant inducible vigor in the wt, *gp1-1*, *act1-5 gp1-1*, and *act1-5* plants. Images were taken at 11 DAT. [F] Total leaf area per seedling (TLA) was quantified for WT, *gp1-1*, *act1-5 gp1-1*, and *act1-5* at 11 DAT. Mean ± SD (n=7). [G] Exogenous application of glycerol decreases plant inducible vigor. Images were taken at 9 DAT. [H] TLA quantification of WT plants with or without glycerol treatment. Mean ± SD (n=9). Fold changes are shown above each group.

### Disruption of GP1-dependent oleic acid homeostasis impairs plant inducible vigor

At2g43710 encodes a stearoyl-ACP desaturase known as SSI2 (Suppressor of SA Insensitive 2), which catalyzes the desaturation of stearic acid (18:0) to oleic acid (18:1) (*21*). Interestingly, transcript levels of *GP1*/*SSI2* in wt plants were increased by GMVs at 1, 2, and 3 DAT (Figure S4A). Such a mild gene induction was not correlated with elevated levels of oleic acid, whereas the *gp1-1* mutant showed decreased levels of oleic acid compared to wt in both mock and GB03 conditions (Figure 2D). Dysfunction of *SSI2* resulted in altered fatty acid composition with increased and decreased accumulation of 18:0 and 18:1, respectively (*21*). The level of 18:1 in the *ssi2* mutant can be restored by a second mutation in the glycerol-3-phosphate (G3P) acyltransferase (ACT1), which catalyzes the acylation of G3P by consuming 18:1 (*22*). To examine whether GP1-dependent plant vigor requires proper homeostasis of 18:1, we generated the double mutant of *act1-5 gp1-1* and examined its responsiveness to GMVs. In the same petri dish, wt plants and the *act1-5* single mutant showed 5.3-fold and 5.4-fold increases, respectively, in plant leaf area; meanwhile *gp1-1* only showed a 1.8-fold increase but the *act1-5 gp1-1* double mutant showed a 3.5-fold increase triggered by the same GMVs (Figure 2E and F). These results suggest that maintaining a proper level of 18:1 is necessary for GMV-induced plant vigor.

To confirm the importance of 18:1 to plant inducible vigor, we treated wt plants with glycerol, which increases ACT1-dependent consumption of 18:1 (*23*). As a result, the glycerol application led to dosage-dependent decreases in GMV-induced plant vigor (Figure 2G-H). Dysfunction of SSI2 or exogenous application of glycerol both induced accumulation of nitric oxide (NO), which is an important regulator of plant development and stress responses including SA-dependent defense (*24*). Consistently, exogenous application of NO to wt plants mimicked *gp1*/*ssi2* mutations in showing impaired plant growth-promotion (Figure S4B-C), indicating that NO-mediated signaling antagonizes GMV-induced plant vigor. These results collectively revealed that GP1/SSI2-dependent 18:1 homeostasis is crucial for plant inducible vigor.

### SA-dependent defense responses partially suppresses GMV-induced plant vigor

Because the *ssi2* mutation causes elevated accumulation of SA and activated SA signaling (*21*), we tested the possible role of SA in the impaired growth promotion phenotype in *gp1* mutant. Firstly, exogenous SA was applied to wt plants. When exposed to GMVs, SA-treated plants showed less increases in fresh weight compared to plants without SA treatments (Figure S5A and B), indicating that the activated SA signaling contributes to the suppression of plant inducible vigor in the *gp1* mutant.

Arabidopsis EDS1 (Enhanced Disease Susceptibility 1) is required for SA production and *R* gene mediated disease resistance (*25*). To investigate whether SA-mediated defense signaling is necessary for the suppression of inducible vigor in *gp1*, we generated an *eds1 gp1-1* double mutant (Figure S5C), and treated the *eds1 gp1-1* double mutant with GMVs. The wt and *gp1-1* showed 4.7 - and 1.7 - fold increases, respectively, in total leaf area; meanwhile the *eds1 gp1-1* double mutant showed a 2.8 - fold increase, thus displaying a partial but significant recovery of GMV-induced plant growth-promotion (Figure 3A and B). Similarly, a partial but significant recovery in the double mutant was also observed when plant freah weight were measured (Figure S5D). Because SA accumulation in plants can be blocked by transgenic expression of *NahG*, which is a bacteria gene that encodes an SA-degrading enzyme (*26*), we also introduced *NahG* into the *gp1-1* mutant. When treated with GMVs, the *NahG gp1-1* plants showed a 3.0 - fold increase in total leaf area, while the growth-promotion in wt and *gp1-1* was 5.2 - fold and 2.0 - fold, respectively (Figure 3C and D). Similarly, measurements of fresh weight showed that *NahG gp1* had a 4.4 - fold increase, while wt and *gp1-1* showed 6.6 - and 3.1 - fold increases, respectively (Figure S5E). Thus it appears that the suppression of inducible vigor in *gp1-1* is only partially due to the activation of SA-dependent defense signaling.

**Figure 3.**
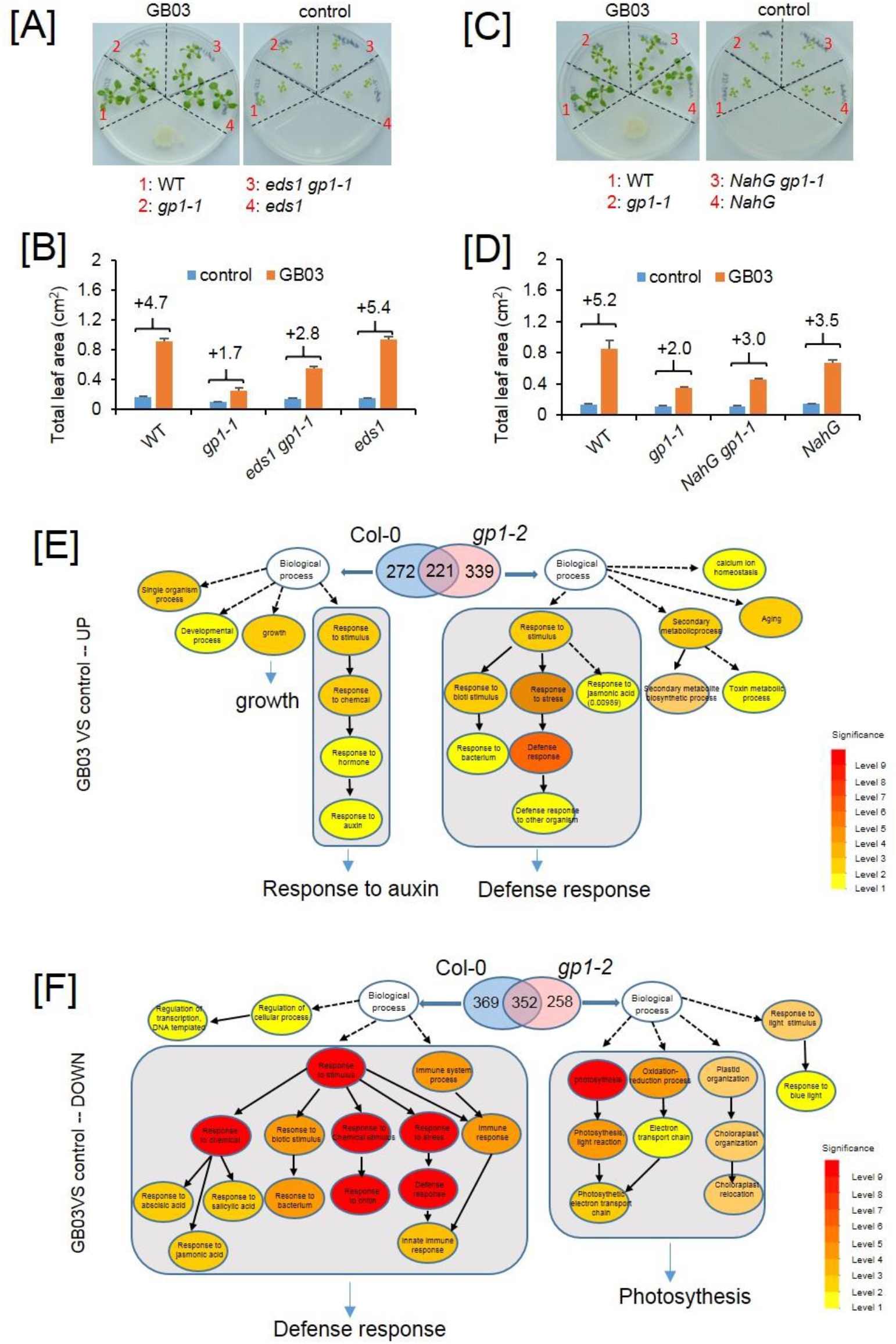
The reduction of plant inducible vigor in *gp1* highlights the anti-correlation between growth and defense. **[A]** Mutation of *EDS1* in *gp1-1* partially restored plant inducible vigor. Images were taken at 10 DAT. **[B]** Total leaf area per seedling was quantified for WT, *gp1-1*, *eds1 gp1-1*, and *eds1* at 10 DAT. Mean ± SD (n=8). **[C]** Transgenic expression of *NahG* in *gp1-1* partially restored plant inducible vigor. Images were taken at 10 DAT. **[D]** Total leaf area (TLA) per seedling was quantified for WT, *gp1-1*, *NahG gp1-1*, and *NahG* at 10 DAT. Mean ± SD (n=8). Fold changes are shown above each group. **[E] and [F]** Gene Ontology (GO) comparative analysis of *Arabidopsis* genes that were induced **[E]** or repressed **[F]** at 2 DAT by GMVs. The blue and pink Venn diagrams show DEGs (differentially regulated genes) identified in Col-0 and *gp1-2*, respectively. The GO pathways are drawn according to AgriGO V2 (http://systemsbiology.cau.edu.cn/). The color key indicates the significance level, in which level 9 means the most significant according to p value of the enrichment. The detailed DEG lists and GO terms are provided in Dataset S1.

### Transcriptome comparison between gp1 and Col-0 highlights the anti-correlation between growth and defense

To further understand how *GP1* dysfunction suppresses plant inducible vigor, we performed transcriptome analyses by RNAseq. The T-DNA insertion line *gp1-2* was compared with its wild type Col-0 under the control and GMV-treated conditions. Totally 493 and 560 differentially expressed genes (DEGs; fold changes ≥ 2, FDR < 0.05) were identified to be up-regulated by GMVs in Col-0 and *gp1-2*, respectively (Figure 3E; Dataset S1). The GMV-up-regulated DEGs in Col-0 showed a GO enrichment of several growth-regulating processes such as auxin responses and cell wall loosening (Figure S6A and B; Dataset S1). Besides, the ferrous iron transporter IRT1 and the ferric iron reductase FRO2, which are crucial for root iron uptake and consequently for photosynthesis (*19*), were also up-regulated (Figure S6C; Dataset S1). These patterns are consistent with previous reports that GMVs-induced plant vigor involves regulation of auxin homeostasis, cell expansion and acquisition of iron (*17, 19*). Gene ontology (GO) analyses were then performed with up-regulated DEGs that were uniquely identified either in Col-0 (272 DEGs) or *gp1-2* (339 DEGs) (Dataset S1). As a result, the up-regulated DEGs in Col-0 showed a GO enrichment in growth related process including auxin response (Figure 3E; Dataset S1). In contrast, the up-regulated DEGs in *gp1-2* not only failed to show GO enrichment of those growth-regulating processes but also were enriched in plant defense responses (Figure 3E; Dataset S1).

In addition to the up-regulated DEGs, GMVs also down-regulated 721 and 610 DEGs in Col-0 and *gp1-2*, respectively (Figure 3F; Dataset S1). The down-regulated DEGs that were unique to Col-0 were enriched with defense response genes (Figure 3F; Dataset S1), whereas the down-regulated DEGs that were unique to *gp1-2* showed GO enrichment of photosynthesis-related genes (Figure 3F; Dataset S1). Therefore, GP1 dysfunction drastically alters plant responses to GMVs by switching the growth-promoting activities to the growth-inhibiting defense responses, resulting in suppression of GMV-induced plant vigor.

### Plants without functional GP1 become defensive to several non-pathogenic microbial factors

Because GMVs decrease defense responses in Col-0 plants (Figure 3F), it is apparent that these microbial products are not considered by wild type plants as pathogen signals. This pattern possibly reflects plant’s ability in distinguishing beneficial microbes from pathogens. On the other hand, the transcriptome comparison between *gp1-2* and Col-0 highlights a group of 52 defense response genes, which are induced by GMVs only in *gp1-2* but not in Col-0 (Dataset S1). This pattern indicates that plants without functional GP1 are no longer able to distinguish GMVs from pathogen triggers. As a result, unlike the wild type plants, *gp1* mutants responded to GMVs with sharp increases in transcript levels of defense-related genes such as the R genes (Figure 4A), as if GMVs were from pathogens even though these microbial products did not cause any observable pathogenic effects on the plants. We compared gene expression levels of *PR1* in the wt, *gp1-1* mutant and the complementation line Pro_GP1_::GP1-FLAG with or without exposure to GMVs. At the time of 2, 3 and 4 DAT, both control and GMV-treated wt plants similarly showed low levels of *PR1* gene expression (Figure 4B). Compared to wt plants, the *gp1-1* mutant exhibited constitutively activated *PR1* gene expression under the control condition; in addition, GMVs strongly elevated mRNA accumulation levels of *PR1* in the *gp1-1* mutant (Figure 4B). Meanwhile, the Pro_GP1_::GP1-FLAG complementation line showed a similar pattern of *PR1* gene expression as that in the wt plants (Figure 4B). Together these observations indicated that wild type Arabidopsis harbor latent defense response (LDR), instead of being inert, to non-pathogenic microbial factors such as GMVs, and that GP1 is essential for Arabidopsis compatibility with *B. amyloliquefaciens* GB03, at least partially through its suppression of plant LDR to GMVs.

**Figure 4.**
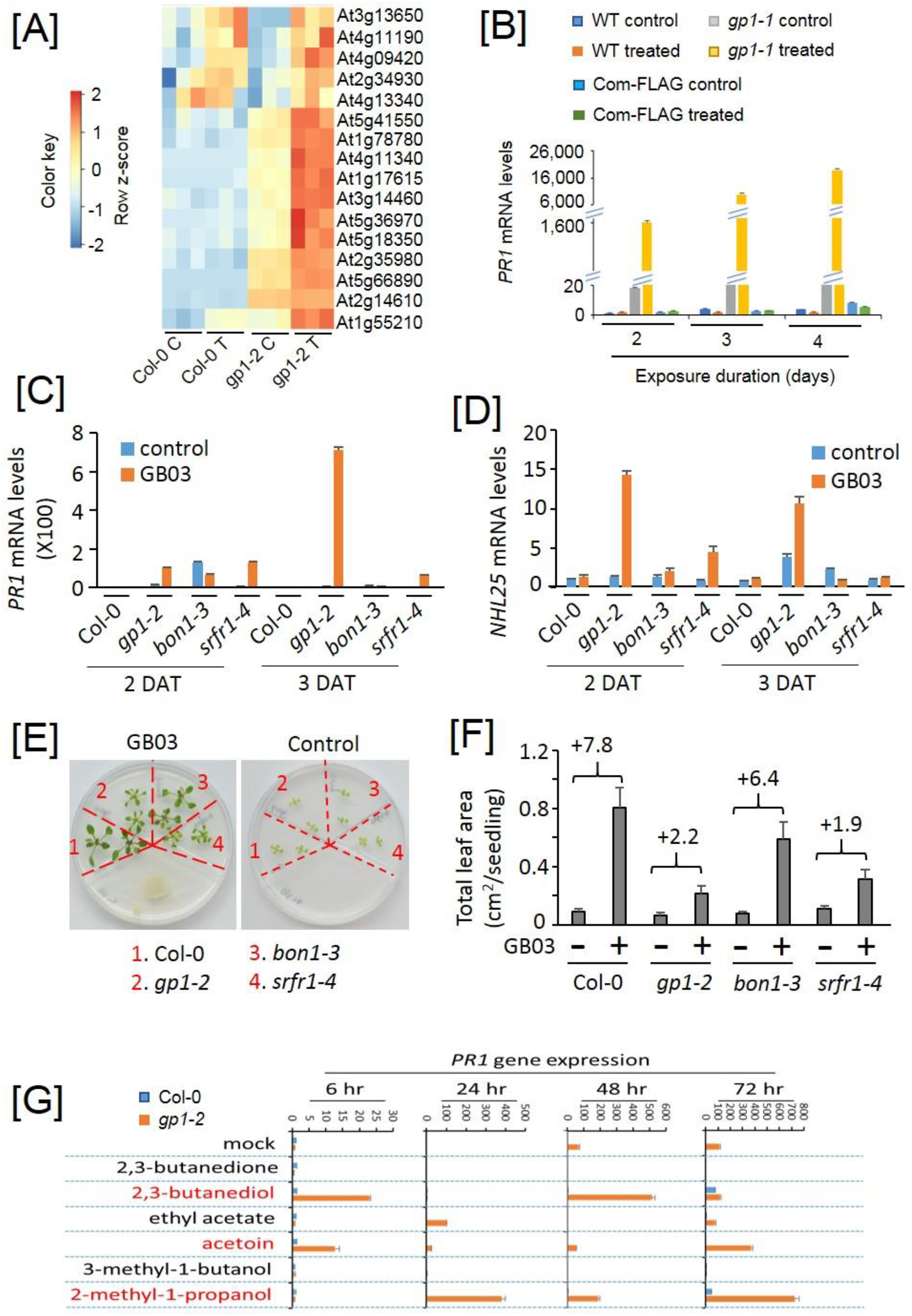
GP1 dysfunction discloses plant latent defense response to non-pathogenic microbial factors. **[A]** A heatmap that shows the expression levels of *R* genes, which were identified as DEGs induced by GMVs only in *gp1-2* but not in Col-0. **[B]** RT-qPCR analysis of Arabidopsis *PR1* gene in WT, *gp1-1* and Pro_GP1_::GP1-FLAG with or without GMVs treatment for 2, 3, 4 days. Values are normalized to *PR1* expression level in the WT control plants at 2 DAT. Mean ± SE (n=3). **[C]** and **[D]** RT-qPCR Arabidopsis *PR1* and *NHL25* genes, respectively, in Col-0 and the *gp1-2*, *srfr1-4*, *bon1-3* mutants with or without GMVs treatment for 2 and 3 days. Values are normalized to *PR1* expression level in Col-0 control plants at 2 DAT. Mean ±SD (n=3). **[E]** Comparison of GMV-induced plant vigor in wild type (Col-0) and the mutants *gp1-2*, *srfr1-4*, and *bon1-3*. Images were taken at 10 DAT. **[F]** Total leaf area quantification of plants with or without exposure to GMVs. Mean ± SD (n=14). Fold changes are shown above each group. **[G]** RT-qPCR analysis of Arabidopsis *PR1* gene in Col-0 and *gp1-2* in response to six individual GMV components. The synthetic compounds were applied at dosages that, when the compounds totally evaporate from the agar-containing solid droplets, would yield in volatile concentrations of 32.5 μg (2,3-butanediol), 7.8 μg (2-methyl-1-propanol), 2.5 μg (3-methyl-1-butanol), 6.2 μg (ethyl acetate), 9.7 μg (2,3-butanedione), and 28.5 μg (acetoin) per mL free space in the petri dish, which resembled the ratio among the six GMV components in natural GMVs as previously reported [33]. Values are normalized to *PR1* expression level in Col-0 mock plants for each time point. Mean ± SD (n=3).

In addition to disclosing LDR to GMVs, dysfunction of GP1 also causes autoimmunity, which is known as elevated defense responses in plants that are not treated with microbes or microbial factors. To investigate the relationship between LDR and autoimmunity, we investigated plant responses to GMVs in *srfr1-4* and *bon1-3*, which are both known as autoimmunity mutants (*27, 28*). The GMV-induced LDR was observed in *srfr1-4* seedlings, although to a less degree compared to that in the *gp1-2* mutant (Figure 4C, 4D). In contrast, *bon1-3* showed no LDR in response to GMVs under the same conditions as *gp1-2* and *srfr1-4* (Figure 4C, 4D). Therefore, although the *gp1* mutants simultaneously display autoimmunity and LDR, autoimmunity does not seem to be a prerequisite for disclosing LDR. Compared to the wild type plants, drastic reductions in GMV-induced plant growth-promotion were observed in *gp1-2* and *srfr1-4*, whereas *bon1-3* displayed only a minor reduction (Figure 4E, 4F). These results further support the anti-correlation between GMV-induced plant growth and defense, since *bon1-3* showed only autoimmunity while *gp1-2* and *srfr1-4* showed both autoimmunity and LDR.

GMVs include over 30 compounds as detected by chromatography coupled with mass spectrometry (GC-MS), with varying individual abundance ranging from 0.6 ng per 24 hr to 614 μg per 24 hr (*29*). In searching for the volatile component(s) that triggered LDR in *gp1*, we examined 6 synthetic compounds individually including 2,3-butanediol, 2-methyl-1-propanol, 3-methyl-1-butanol, ethyl acetate, 2,3-butanedione and acetoin, which are the major components of GMVs and are commercially available. Agar-containing solid droplets were used to carry the compounds for progressive release of the volatiles. By comparison with Col-0, *gp1-2* displayed LDR to 2,3-butanediol, acetoin, and 2-methyl-1-propanol, as shown by the hyper induction of *PR1* gene expression at 6, 24, 48 and 72 hrs after the chemical treatments (Figure 4G). In contrast, 2,3-butanedione and 3-methyl-1-butanol did not induce *PR1* gene expression at the examined time points, whereas ethyl acetate transiently induced *PR1* expression in *gp1* only at 24 hr after the treatment (Figure 4G). Together with the observations of GMV-treated plants, the results of synthetic compounds demonstrate that Arabidopsis *GP1* dysfunction leads to the LDR in plants exposed to the beneficial rhizobacterium GB03.

### GP1 dysfunction alters the assemblage of root-associated rhizobacteria community

To further explore the role of GP1 in plant interactions with microbes, we investigated the rhizosphere and root endosphere microbiomes of *gp1-1*, wt, and the Pro_GP1_::GP1-FLAG complementation line. Because the wt plant is in the Col-0 background and contains a *35S::SUC2* transgene that encodes a sucrose transporter (*30*), the microbiomes associated with Col-0 were examined in parallel so as to further assess potential impacts of *35S::SUC2* on the root microbiome. The bacteria communities were profiled by 16s rDNA sequencing of DNA samples from the three compartments of bulk soil, rhizosphere, and endosphere. A total of 8432516 high quality sequences were obtained with a median read count per sample of 168312.5 (range 22843-690623) across all the 46 samples. To correct the sequencing depth differences among samples, these high quality sequences were further subjected to rarefaction to 18000 sequences per sample based on the rarefaction curve (Figure S7A), which resulted in 8030 unique OTUs as the threshold-independent community (TIC) (Dataset S2). A minimum of 20 sequences per OTU in at least one sample was used as a criterion to define Abundant Community members (ACM) (*31, 32*). The ACM of all samples was represented by 572 bacterial OTUs comprising of 73.8% of the rarefaction quality sequences (Dataset S2). We normalized this table by dividing the reads per OTU in a sample by the sum of the usable reads in that sample, resulting in a table of relative abundances (RA) (Dataset S2) for quantitative comparison.

By using weighted UniFrac metric on ACM (*33*), we examined the main factors accounting for the community diversification across the samples. The hierarchical clustering of UniFrac distances clearly separated the bulk soil samples from the rhizosphere and the endosphere samples, indicating that the compartment is a major determinant of bacteria communities (Figure 5A). Analyses of taxonomic distribution showed that the bulk soil community mainly consisted of the phyla of Proteobacteria, Bacteroidetes, Acidobacteria and Actinobacteria, whereas the rhizosphere and the endosphere communities were dominated by Proteobacteria and Bacteroidetes (Figure S7B).

**Figure 5.**
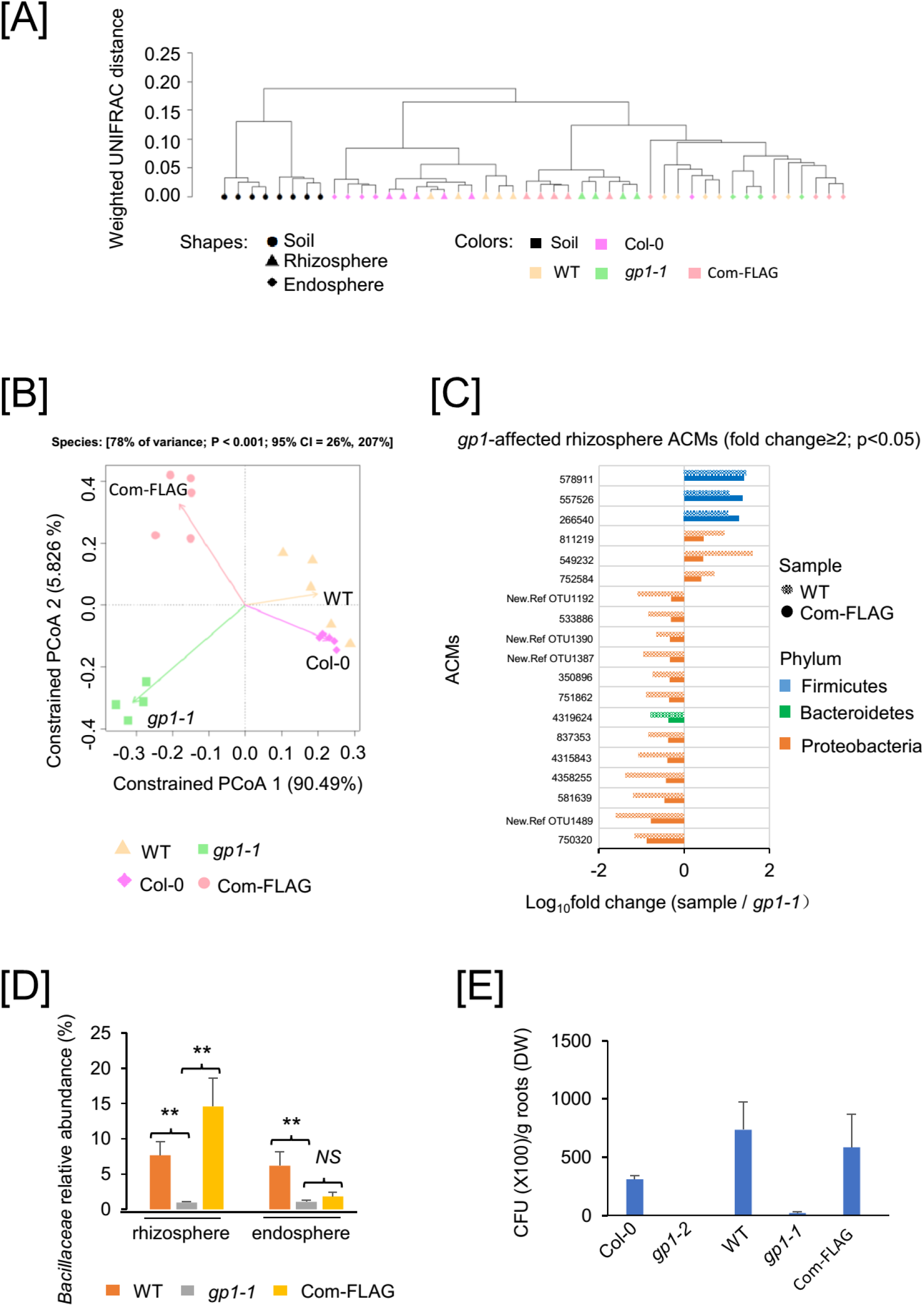
GP1 dysfunction alters the assemblage of root-associated rhizobacteria community. **[A]** Beta diversity of the Abundant Community members (ACM). Between-sample similarities were estimated on 18 000 sequences per sample using the phylogeny-based UniFrac distance metric. Weighted UniFrac is sensitive to the sequence abundances. **[B]** Constrained principal coordinate analysis (PCoA) of ACM in the rhizosphere of WT, *gp1-1*, Col-0 and Pro_GP1_::GP1-FLAG. The percentage of variation explained by each axis refers to the fraction of the total variance of the data (ACM) explained by the constrained factor. **[C]** ACMs that are significantly changed in *gp1-1* mutant and complemented by Pro_GP1_::GP1-FLAG in the rhizosphere compartment, as determined by fold changes of ACM abundance in WT and Pro_GP1_::GP1-FLAG versus *gp1-1* (RA≥5‰; Fold change ≥ 2, Tukey p<0.05). **[D]** Relative abundance of the *Bacillaceae* family in the examined rhizosphere and endosphere compartments. ** Tukey, P < 0.01; NS, not significant; **[E]** Root colonization of GB03 in Col-0, *gp1-2*, WT, *gp1-1* and Pro_GP1_::GP1-FLAG plants. Mean ± SE (n=4).

**Figure 6.**
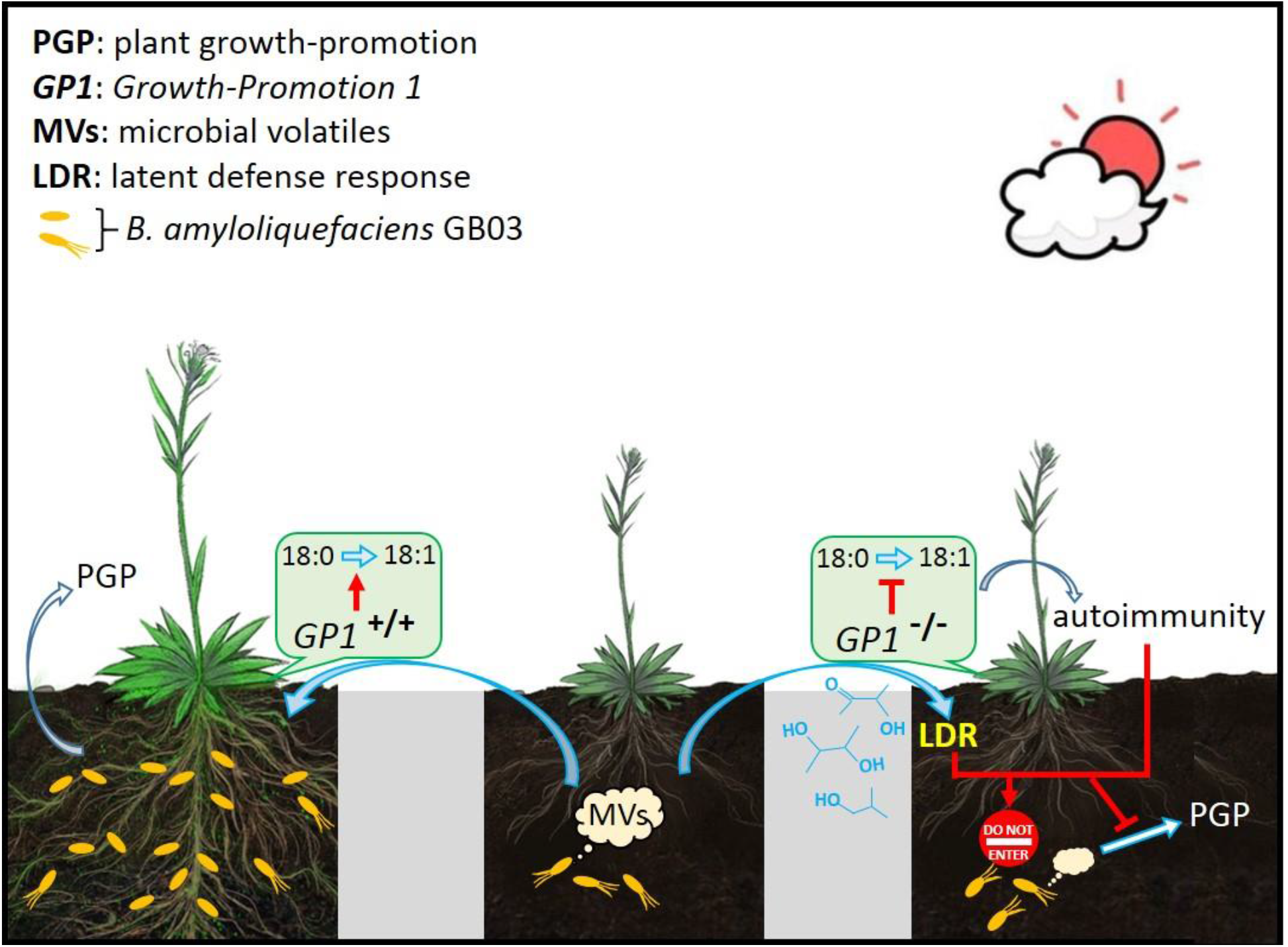
A model depicting the role of GP1 in Arabidopsis interaction with *B. amyloliquefaciens* GB03. GP1 catalyzes the desaturation of stearic acid (18:0) to oleic acid (18:1) and is required for plant growth-promotion triggered by the beneficial rhizobacterium GB03. Dysfunction of GP1 not only causes autoimmunity in the absence of bacterial exposure, but also discloses latent defense response (LDR) in response to GB03-produced volatile compounds that, in contrast, are non-pathogenic to wild type plants. Consequently the activated plant defense in *gp1* represses GB03-induced plant growth-promotion. In addition, *gp1* shows altered assemblage of the root-associated rhizobacteria community as well as severely impaired root colonization of GB03. The chemical structures of acetoin, 2,3-butanediol, and 2-methyl-1-propanol, which are three microbial volatile compounds identified as elicitors of LDR, are shown in blue from the top to the bottom.

We next performed constrained principal coordinate analysis (PCoA) of ACM to evaluate the impacts of GP1 dysfunction on bacteria communities in the rhizosphere and the endosphere. The PCoA plots clearly separate *gp1-1* from wt along the major coordinates, which represent 90.49% and 77.48% variances for the rhizosphere and the endosphere, respectively (Figure 5B; Figure S8A), indicating that GP1 dysfunction alters the microbial communities in these two compartments. The PCoA plots also clearly distinguish *gp1-1* from Pro_GP1_::GP1-FLAG; however, transgenic expression of Pro_GP1_::GP1-FLAG in *gp1-1* resulted in only partial restoration of the bacteria communities associated with wt roots (Figure 5B; Figure S8A). Because the Pro_GP1_::GP1-FLAG complementation line displayed fully-recovered growth phenotype of GMV-inducible vigor (Figure 2B-C), we deduced that *gp1-1* may contain additional EMS-generated mutations, which caused GP1-independent alterations in root-associated microbiomes. Notably, rhizosphere abundance of the phyla Verrucomicrobia and Firmicutes was increased (Tukey, p=0.00027) and decreased (Tukey, p=0.00022), respectively, in *gp1-1* compared to the wt; whereas in Pro_GP1_::GP1-FLAG, both Verrucomicrobia (Tukey, p=0.045) and Firmicutes (Tukey, p=0.00144) showed significant restoration compared to *gp1-1* (Figure S8B; Dataset S2), indicating that GP1 has strong influence on the association between these two phyla and Arabidopsis. Similarly, endosphere abundance of the phyla Planctomycetes and Verrucomicrobia was dependent on GP1, as demonstrated by comparisons of the *gp1-1* with wt and with Pro_GP1_::GP1-FLAG (Figure S8C; Dataset S2). Together these results revealed an influential role of GP1 in the assembly of plant-associated bacteria community.

*B. amyloliquefaciens* GB03 belongs to the family of *Bacillaceae* within the phylum of *Fimicutes*. Among the ACMs that are significantly changed in *gp1-1* mutant and complemented by Pro_GP1_::GP1-FLAG (RA≥5‰; Tukey, p<0.05) in rhizosphere compartment, those belonging to the phylum of Firmicutes are all decreased in *gp1-1* mutant (Figure 5C, Dataset S2). However, in the endosphere compartment, none of the complemented ACMs are from Firmicutes (Figure S8D; Dataset S2). At the family level, rhizosphere abundance of *Bacillaceae* in *gp1-1* was markedly reduced (Tukey, p=0.0011) compared to the wt; and it was restored by transgenic expression of Pro_GP1_::GP1-FLAG (Figure 5D; Dataset S2). In contrast, although endosphere abundance of *Bacillaceae* was also decreased by GP1 dysfunction, it was not restored by the expression of Pro_GP1_::GP1-FLAG (Figure 5D). Thus it appears that GP1 particularly affects plant association with those *Bacillaceae* members that dwell in the rhizosphere instead of the endosphere. We next examined whether GP1 dysfunction affects GB03 colonization. Compared to their wild type plants, the *gp1* mutants showed drastic reductions in the abundance of root surface-associated GB03 (Figure 5E); meanwhile GB03 colonization to roots was restored in the Pro_GP1_::GP1-FLAG plants (Figure 5E). Therefore, the loss of inducible vigor in GB03-innoculated *gp1* plants in the soil can also be attributed to the impaired bacteria colonization, in addition to the mistakenly activated defense responses that would directly compromise growth.

## Discussion

Plants have evolved to distinguish beneficial microbes from pathogens, leading to either mutualistic association or activated immune responses for optimized survival (*8–10*). Precise recognition of microbial factors is critical for plant-microbe interactions. In this study, we demonstrate that dysfunction of Arabidopsis GP1 causes the plant to become defensive in response to the beneficial rhizobacterium *B. amyloliquefaciens* GB03 and consequently lose GB03-triggered plant growth-promotion. The microbial volatiles released from GB03 are non-pathogenic to wild type plants, whereas the same microbial factors trigger plant defense responses in the *gp1* mutants. This misrecognition discloses a hidden layer of plant innate immunity, which is proposed herein as latent defense response (LDR).

LDR refers to host defensive responses elicited by non-pathogenic microbial factors. Plant compatibility with commensal or beneficial microbes requires a seemingly inert relationship between non-pathogenic microbial factors and the host. The inertness can be either truly irresponsiveness or due to suppression of plant innate immunity. In addition to microbe-mediated suppression of plant immunity for establishing mutualism (*10, 20*), the discovery of LDR caused by GP1 dysfunction has demonstrated that suppression of plant immunity for establishing commensalism or mutualism can be on plant’s own initiative, that is, when plants show no defensive response to certain non-pathogenic microbial factors, it does not mean that these microbial factors are really inert to plants, instead, it can be because plants are being regulated by certain internal factors so that they would not respond to commensal or beneficial microbes in a hostile way.

LDR is different from autoimmunity, because the former is elicited by non-pathogenic microbial factors, whereas the latter refers to an elevated level of defense without needing any exposure to microbes or microbial factors (*34*). LDR is also different from induced systemic resistance (ISR), which refers to the priming of plant resistance to pathogens as a result of mutualistic association between plants and certain PGPR (*35, 36*), because LDR leads to plant incompatibility with beneficial microbes, whereas ISR acts against pathogens without impairing the plant-PGPR association. Although *gp1* displayed LDR in response to at least three GMV components, LDR in this mutant was not elicited by 2,3-butanedione that is also known as diacetyl. We recently reported that diacetyl triggers integrated modulation of plant immunity and phosphate (Pi) starvation responses, resulting in opposing relations between Arabidopsis and *B. amyloliquefaciens* GB03 depending on plant Pi availability (*20*). Particularly, while diacetyl facilitates mutualistic association between Pi-sufficient Arabidopsis and GB03 through suppression of plant immunity, defense responses are strongly induced by diacetyl in Pi-deficient plants. These results suggest that diacetyl is an elicitor of plant LDR that depends on Pi availability. Therefore, GP1-dependent LDR and Pi-dependent LDR jointly highlight the complexity of the yet-to-be-explored regulatory network of plant LDR.

GMV-induced plant vigor is contributed by several growth-promoting processes including regulation of auxin homeostasis (*15, 17*). Consistent with the negative impact of GP1 dysfunction on plant inducible vigor, transcriptome analysis of *gp1-2* revealed a loss of GMV-dependent regulation of auxin hemostasis. This observation is consistent with a previous report that GP1 dysfunction repressed transcript levels of auxin signaling-associated genes (*37*). It should be noted that the bacterial volatile emission contains not only organic components but also carbon dioxide (CO_2_) that is necessary for plant photosynthesis. Elevated CO_2_ levels alone in sealed plates can also increase plant growth (*38*). However, having the same phenotype should not lead to the deduction that one trigger (e.g., bacterial volatile emissions) necessarily functions through the same way as the other (e.g., CO_2_). In fact, example evidence has been shown that the fertilization effects of CO_2_ are not mediated through the same mechanisms as bacterial volatile emissions (*39*). Because CO_2_ is essential for plant growth, it is likely that bacteria volatile-induced plant growth-promotion reflects the combined effects from both the specialized organic components and CO_2_. Nonetheless, that GP1 dysfunction impairs GB03-induced plant vigor was demonstrated not only in sealed plates, but also in soil pots where direct bacteria-plant interaction occurred. More importantly, GMV-induced plant LDR in *gp1* were further supported by the treatments with individual synthetic compounds.

GP1 and its S-ACP-DES isozymes are conserved in diverse plant species including Arabidopsis, soybean, wheat and rice (*40, 41*). Arabidopsis GP1 dysfunction not only reduces root colonization of *B. amyloliquefaciens* GB03, but also decreases the abundance of root-associated *Bacillaceae* family, which is known to contain many plant growth-promoting rhizobacteria species (*4*). In addition to *Bacillaceae*, GP1 dysfunction also affects root-association of bacteria belonging to several other families. Since plant microbiota can be influential to host fitness (*1, 8*), it is conceivable that GP1 dysfunction may have broader impacts than plant-microbe interactions, as a result of indirect regulation through affecting root-associated microbiota.

## Materials and Methods

### Plant and bacteria materials and growth conditions

The *gp1-1* mutant was isolated from an EMS mutagenized library generated previously, which are in Col-0 background and contains a *35S::SUC2* transgene (*30*). The *gp1-2* (SAIL_209_D07), *act1-5* (SALK_069657) and *eds1* (SALK_057149) mutants were ordered from the NASC (Nottingham Arabidopsis Stock Centre) or ABRC (Arabidopsis Biological Resource Center). *NahG* was from Prof. Alberto Macho in Shanghai Center for Plant Stress Biology (PSC). The double mutants of *act1-5 gp1-1*, *eds1 gp1-1* and *NahG gp1-1* were generated by genetic crosses between *gp1-1* and *act1-5*, *eds1*, and *NahG*, respectively. The *srfr1-4* (Sail_412_E08) and *bon1-3* (SALK_200380) was from Prof. Yang Zhao at PSC.

For plant growth conditions, the wt, *gp1-1*, *act1-5 gp1-1*, *eds1 gp1-1* and *NahG gp1-1* seeds were surface sterilized using 30% bleach, washed and sowed on ½ MS containing 1% glucose and 0.7% agar (pH5.7). After stratification at 4°C for two days, plates were moved to a growth chamber at 22°C with 100μmol/m^2^/s fluorescent light (16 h of light and 8 h of dark).

The GB03 (*Bacillus amyloliquefaciens*) from the lab stock was cultured in LB liquid medium at 37°C overnight (OD 600=1.0-1.2) with shaking at 220 rpm before use.

### Bacteria treatment in VOC system

For the GB03 VOC treatment, the Y-plates were used which have 3 partitions inside, two of which are for seedlings and another is for bacteria treatment. 5-day-old seedlings were transferred to new half MS medium in two partitions of a Y-plate, while 20 μl of the bacteria GB03 was added into another partition. By this way the bacteria will not touch the plants but its emitted VOCs can spread inside and affect the plant. Control was performed by adding no bacteria. The plates were then sealed with parafilm and placed at 22°C, 16h light/8h dark condition. Photos were taken when the growth promotion phenotype of Col-0 were clear (Usually at 9-11 DAT). The pictures were imported into the software Image J, and the plate diameter was used to calibrate the total leaf area.

### Bacteria treatment in soil system

For the GB03 treatment in soil system, the soil was mixed with vermiculate with a ratio of 2:1 (soil: vermiculate), filtered with a filter sieve, autoclaved at 121 °C for 12 h, and wetted with water containing 0.1% green fertilizer. The soil was then evenly distributed into pots, and 5-day-old seedlings were transferred into the square pots and grown in a growth room at 22°C for another day. For the bacteria treatment, 10 μl of the GB03 stock was pre-inoculated in 5 ml LB at 37°C overnight. Then the 5 ml bacteria culture was added into 150 ml LB and shaking at 37°C for another 6 h (OD600=1). Centrifuge at 4000 rpm for 25 min and discard the supernatant. The bacteria precipitation was re-suspended in equal volume of 0.45% sterilized NaCl solution. Add 5 ml of the bacteria suspension to the rhizosphere of each plant. For control, only 0.45% NaCl was added. Photos were taken 9 days after treatment. Leaf area and fresh weight were measured 11 days after treatment.

### Bacteria treatment in soil-in-tube system

For bacteria treatment in soil-in-tube system tube, the soil was mixed with vermiculate with a ratio of 2:1 (soil: vermiculate), filtered with a filter sieve, autoclaved at 121°C for 12 h, and wetted with water. 3 grams of the soil was weighed, filled in a Nonwoven cloth package, and fixed in a 50 ml tube. For each tube, a seedling of 5 days grown on half MS was transferred to the soil and watered with 2 ml of water containing 1 ‰ fertilizer. 25 μl GB03 bacteria culture was inoculated on a small cap filled with half MS solid medium and cultured overnight. The small cap with the GB03 bacteria was then put in the bottom of the 50 ml Tube to release the VOC while not touch the plant. Control was performed by putting half MS without bacteria but with kanamycine and ampicillin to avoid contamination. The plants were watered with 1 ml water per day for each seedling. 14 days after treatment, photos were taken and the fresh weight of the aerial part was weighed.

### EMS mutant screening and map based cloning

Ethyl methanesulfonate (EMS)-mutagenized M2 seeds (*30*) were grown on half MS medium containing 1% glucose and 0.7% agar in Y-plates. After GB03 treatment for 7 days, the wt plants show obvious growth promotion. Putative mutants with reduced growth promotion were transferred to soil. The phenotype was then confirmed using the newly harvested seeds. The mutants with confirmed phenotype was used for further study.

For map-based cloning, *gp1-1* mutant in Col-0 ecotype was crossed to a wt plant in Ler ecotype. The resulting F1 was selfed to get the F2 seeds. The F2 progenies were subjected to GB03 treatment, and the individuals which show reduced GB03-mediated growth promotion phenotype as *gp1-1* were selected as the F2 mapping population. Using 214 F2 individuals, the gene was mapped to Chr2 and narrowed down to an interval of 175K between 17 970 K to 18 145K. Then the genomic DNA of *gp1-1* mutant was extracted and whole genome was sequenced at the Core Facilities of Genomics of Shanghai Center for Plant Stress Biology. The resulting sequence was than aligned with the Col-0 reference to search for the mutations.

### Complementation test

For complementation test, the genomic sequence including 2kb upstream the At2g43710 gene was amplified by PCR using the Col-0 genomic DNA as template. Primers are listed in Table S1. The PCR product was then inserted into the plant expression vector PCAMBIA1305-3HA, PCAMBIA1305-3MYC and PCAMBIA1305-3FLAG, respectively, between the restriction site EcoR1 and Xba1. The resulting constructs were then introduced to *gp1-1* mutant by the agrobacteria-mediated transformation. The homozygous T3 transgenic lines were used for the further experiments.

### Oleic acid analysis

Oleic acid contents measurement was performed as described previously (*42*) with modifications. For details, 5-day-old seedlings grown on half MS medium were treated with GMVs for 3 days before the seedlings were harvested. 20 mg of the fresh plant materials were placed in a 2.0 mL tube, frozen in liquid nitrogen and homogenized. Samples were then submitted for oleic acid analysis in the Core Facility of Plant Proteomics and Metabolomics in Shanghai Center for Plant Stress Biology, China. Briefly, 0.9 mL of 1:2 chloroform: methanol (v/v) mixture containing 0.05 μmol heptadecanoic acid was added into the tube containing the sample. The tube was then vortexed at 120 rpm and 35 °C for 1 h. After that, 0.3 mL of chloroform was added to the tube and the tube was vortexed at 35 °C for 5 min. Then 0.5 mL H 2O was added to the tube and the tube was vortexed at 35 °C for 5 min. The tube was centrifuged at 5000 rcf for 10 min at 4 °C. Then, 0.5 mL solution from the down phase was collected and transferred into a 1.5 mL tube. The solution was dried under vacuum at 4 °C for 1 h, and 0.5 mL 4% H^2^SO_4_ in MeOH was added in to the tube. The tube was capped and heated at 85 °C for 1 h. After cooling, 0.6 mL n-hexane was added into the tube. The tube was shaken thoroughly and centrifuged at 5000 rcf for 3 min. Then, 0.5 mL solution form n-hexane phase was transferred into a new HPLC vial. The vial was dried under vacuum at 4 °C for 30 min. The sample was re-dissolved in 50 μL n-hexane, and 1 μL solution was injected into GC-MS for analysis.

### RNA seq and analysis

For RNA seq, Col-0 and *gp1-2* (SAIL_209_D07) were grown on ½ MS medium containing 1% sucrose. 5-day-old seedlings were transferred to a Y plate and treated with GB03 for 2 days before the seedlings were harvested. Control was performed by no GB03 treatment. The total RNA was extracted using the Trizol reagent. Each sample has 3 biological replicates. The library constructs and sequencing were performed by the Core Facilities of Genomics in Shanghai Center for Plant Stress Biology.

For data analysis, the raw data is preprocessed by trimmomatic (version 0.36). Then, the clean reads were aligned to the TAIR10 reference genome using hisat2 (version 2.1.0) with parameters “--rna-strandness RF -p 30 --dta”. Reads_count matrix was produced by htseq-count (version 0.9.1). Then we used edgeR to get differential expression genes (DEGs) with the cutoff of " fold change ≥2 and FDR <0.05". The gene ontology (GO) enrichment analysis was performed using the online tool of “agriGO v2.0” (*43*). The heatmaps were generated using the software of pheatmap of R package or the online tool of iDEP.90 (http://bioinformatics.sdstate.edu/idep/).

### Treatments with individual volatile compounds

Arabidopsis seeds were allowed to germinate on ½ MS agar medium in Y-plates, which contained three portions separated by three inner partitions. Seedlings at 5 days-after-germination were treated with individual synthetic compounds as previously reported (*20*). Briefly, the compounds were applied at dosages that, when the compounds totally evaporate from the agar-containing solid droplets, would yield in volatile concentrations of 32.5 μg (2,3-butanediol), 7.8 μg (2-methyl-1-propanol), 2.5 μg (3-methyl-1-butanol), 6.2 μg (ethyl acetate), 9.7 μg (2,3-butanedione), and 28.5 μg (acetoin) per mL free space in the petri dish, which resembled the ratio among these six GMV components in natural GMVs as previously reported (*29*).

### Quantitative Real time PCR

For real time PCR, total RNA was extracted from by Trizol reagent. 1μg of the total RNA was subjected to reverse transcription using the Transcript One-Step gDNA Removal and cDNA Synthesis Super Mix kit (Transgene) in 20 μl volume according to the manufacturer’s instructions. Q-PCR was performed using the iTaq Universal SYBR Green supermix (Biorad) in 20 μl volume using CFX96 real-time PCR detection system (BioRAD). *Actin 2* was used as the internal control. All the primers are listed in Table S1.

### Preparation of metagenomic DNA and 16s rDNA sequencing

The field soil used for microbiota experiments was collected at the Chenshan Botanical Garden, China. The soil was cleaned from plant parts, worms and stones and homogenized manually. The cleaned and homogenized field soil was mixed with the experimental soil in the ratio of 1:1. The seeds of Col-0, wt, *gp1-1*, and Pro_GP1_::GP1-Flag/*gp1-1* were first sowed on ½ MS medium and grown in a growth chamber (22°C, 16h light/ 8h dark). Then the 7 days old seedlings were transferred to pots containing the mixed soil, with the density of 5 seedlings per pot, and grown at growth room (22 °C, 16h light / 8h dark) for another 3 weeks until harvest. Unplanted pots were subjected to the same conditions to prepare the control soil samples. Two pots (10 seedlings) were defined as 1 biological replicate. At least 4 biological replicates were prepared for each genotype. The metagenomic DNA from soil, rhizosphere and root samples were prepared as described (*44*).

Amplicon libraries were generated using the PCR primers 799F (AACMGGATTAGATACCCKG) and 1193R (5’-ACGTCATCCCCACCTTCC-3’) spanning ~400 bp of the hypervariable region V5-V7 of the bacterial 16S rRNA gene (*45, 46*). For details, the first-round PCR was performed to amplify the targeted genomic DNA with primers of 799F-B and 1193R-B with common bridging sequences (5’-ggagtgagtacggtgtgc-3’ and 5’-gagttggatgctggatgg-3’) added at the 5’ end of the 799F and 1193R (see Table S1). The PCRs were assembled in a laminar flow hood in 25 μl volume including the following components: 12.5 μl of 2X KAPA HiFi HotStart ReadyMix, 5 μl of 1 μM of each fusion primer and 2.5 μl of 5 ng/μl adjusted template DNA. The PCR reactions were performed using the condition: 95°C for 3 min, 30 cycles of: 95°C for 30 seconds, 55°C for 30 seconds, 72°C for 30 seconds and extension at 72°C for 5 min. Each reaction has three technical replicates. The resulting PCR products were loaded on a 2% agarose gel and a band of ~400 bp were cut from the gel and extracted using the QIAquick Gel Extraction kit (Qiagen). The DNA concentrations were determined using the Qubit™ dsDNA HS Assay Kit on Qubit®2.0.

The first-round PCR products were further barcoded during the second-round PCR as described previously with modification (*47*). Common primers for the second-round PCR included platform-specific adapter sequence, fixed barcode sequence and bridging sequence. The second amplification was conducted in 20 μL volume containing 10 μl of 2x KAPA HiFi Hot Start Ready Mix, 200 nM of index primers 2P-F and 2P-R, 2 nM of barcode primers of F-(N) and R-(N), and 40ng PCR product (*47*). The PCR was run with the conditions: 95°C for 3 min, 10 cycles of: 95°C for 30 seconds, 55°C for 30 seconds, 72°C for 30 seconds, and extension at 72°C for 2 min. Each reaction has 2 technical replicates. After second PCR, the PCR products were gel-extracted using the QIAquick Gel Extraction kit and eluted in 30 μl ddH 2O. After purification, technical replicates were combined to one biological sample, and DNA concentrations were determined using the Qubi dsDNA HS Assay Kit on Qubit 2.0. Then all the biological samples were mixed with the same amount and sent to the core facility of genomics in Shanghai Plant Stress Biology Center for sequencing.

### Microbiota data analysis

For data analysis, the raw data quality was controlled and preprocessed using FastQC v0.11.8 and trimomatics v0.36 (*48*), and the processed high quality data was assembled with FLASH v1.2.11 (*49*), requiring an overlap of at least 10 bp (-m 10) for the two paired-end reads. For subsequent analysis, we mainly used QIIME v1.91software (*50*). The chimeric sequences were removed using the usearch (-m usearch61) method of the script *identify_chimera_seqs.py* with "Gold" database (-r gold.fa). The remaining high quality sequences were clustered using the script *pick_open_reference_otus.py* and OTUs (operational taxonomic units) were classified. A sequence with more than 97% identity is clustered, and only at least 10 sequences in a cluster were output (-m 10). The OTUs belonging to mitochondria, Chlorophyta, Archea and Cyanobacteria were then removed using the script *filter_taxa_from_otu_table.py* w. By this way, a total of 8432516 high quality sequences were obtained with a median read count of 168312.5 per sample (range: 22843-690623). The high quality reads were clustered into 10061 microbial OTUs.

Next, we used the function *calculateRarefaction* of the R package ShotgunFunctionalizeR (*51*) to evaluate the rarefaction curve. Then we used the script *multiply_rarefaction.py* of QIIME to generate the rarefied tables (100x tables from 18000 sequences per sample, step of 180 sequences) and generated a table composed of 8030 OTUs as the threshold-independent community (TIC) (*31*). Among the TIC, a minimum of 20 sequences per OTU in at least one sample was used as a criterion to define Abundant Community members (ACM) (*31*). The relative abundance (RA) of each ACM in a sample was calculated by dividing the reads of the ACM by the sum of the usable reads in that sample. The relative abundance of each phylum or family was the sum of the ACMs belonging to that phylum or family in a sample. The significant differences between samples were assessed by the ANOVA-based statistics with Tukey’s HSD test. The UniFrac distance was calculated using the script *beta_diversity.py* to evaluate beta-diversity between samples based on the ACM. Heatmaps were performed by the online tool of iDEP.90 (http://bioinformatics.sdstate.edu/idep/). The “*gp1*-affected ACMs” are selected by the following threshold: (1)Relative abundance≥5‰_;_(2) Significantly changed in *gp1-1* mutant compared to wt (wt VS *gp1-1* P<0.05); (3) Restored by the complementation line Pro_GP1_::GP1-FLAG /*gp1-1* (Pro_GP1_::GP1-FLAG /*gp1-1* VS *gp1-1* P<0.05).

### Root colonization test

The seeds of wt, *gp1-1*, Pro_GP1_::GP1-FLAG/*gp1-1*, Col-0 and *gp1-2* were sowed on 1/2 MS containing 1% glucose (for wt, *gp1-1*, Pro_GP1_::GP1/*gp1-1*) or sucrose (for Col-0 and *gp1-2*) and 1% agar, stratificated at 4°C for 2 days and grown vertically in a chamber at 22°C, 16h light / 8h dark. The bacteria root colonization assays followed Morcillo et al. (2020). Briefly, the GB03 was cultured in LB liquid medium at 37 °C overnight (OD =0.8-1.0). The bacteria was collected by centrifuge at 4000rpm for 15min and resuspended in sterile 0.45% NaCl (OD=0.8). Then 10-day-old seedlings were incubated with the resuspended bacteria (10 seedlings in 1ml culture volume) for 24 hours at 22 °C. The roots were collected and washed in sterile water once, sterilized in 70% ethanol for 50s and washed in sterile water for 4 times. After that, the roots were ground using metallic beads and resuspended in 1ml of sterile water, diluted, spread in solid LB medium and incubate at 37 °C overnight. The colony numbers were counted the next day and divided by the dry weight of the roots. The values are mean ± SE (n=4).

## Acknowledgments

We thank Prof. Choong-Min Ryu at Korea Research Institute of Bioscience & Biotechnology for *B. amyloliquefaciens* GB03, Prof. Alberto Macho at Shanghai Center for Plant Stress Biology (PSC) for *NahG*, and Prof. Yang Zhao at PSC for *srfr1-4* and *bon1-3*. We thank the PSC Core Facility of Genomics for sequencing service and the PSC Core Facility of Plant Proteomics and Metabolomics for oleic acid measurement. We also want to make a special thanks to Alejandro Álvarez for his artistic collaboration in the design of the model presented in this manuscript.

## Author contributions

H.Z. designed the project; S.C. and Y.Y. performed all the experiments and data analyses; L.P. performed bioinformatics analyses on RNAseq and microbiota data. X.L., R.K., F.Y., S.K.S., D.H., S.L., J.I.V., R.J.L.M., W.W., W.H. and M.L. participated in the experiments and/or data analyses. H.Z. wrote the manuscript with input from Y.Y., C.P.S., J-K. Z. and P.W.P.

## Data and materials availability

The raw RNAseq data is available in the NCBI GEO with the accession number GSE139154 (token: gjwdyioonnaxnsp); The V5-V7 region of 16 S rRNA gene sequencing data generated during the current study are available in the NCBI SRA under BioProject PRJNA578576.

## Supplementary Materials

Figures S1-S8

Table S1

External Databases S1-S2

