## Supplementary Materials for "Latent defense response to non-pathogenic microbial factors impairs plant-rhizobacteria mutualism"

**This PDF file includes:**

Figs. S1 to S8
Table S1
Captions for Data S1 to S2

**Other Supplementary Materials for this manuscript include the following:**

Data S1:
DEG (differentially expressed gene) lists generated from the RNA seq data

Data S2:
Micobiota datasets

[A]

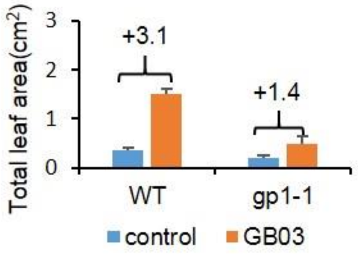


[B]

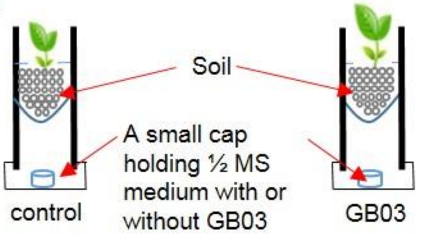


[C]

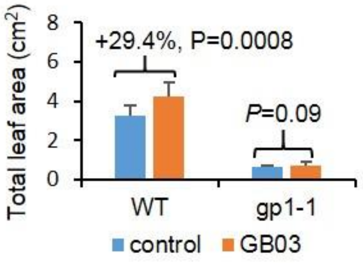


**Figure S1. *gp1* mutation impairs GB03-induced growth promotion in Arabidopsis. [A]** Comparison of *gp1-1* with its wild type (WT) as measured by total leaf area per seedling (TLA). Plants were grown in 1/2-strength MS medium as shown in Figure 1A. Samples were harvested at 11 DAT. Mean ± SD (n=7). **[B]** A diagram showing the experimental setup of the soil-in-tube system. See Materials and Methods for details. **[C]** Comparison of *gp1-1* with its wild type (WT) as measured by total leaf area per seedling (TLA). Mean ± SD (n=7). Plants were grown in soil with or without GB03 inoculation as shown in Figure 1E. Samples were harvested at 11 DAT. Mean ±SD (n=12). Differences between the treated and the control plants were shown as fold changes above each group for all quantification of TLA. Student t-test p values are shown. Related to Figure 1.

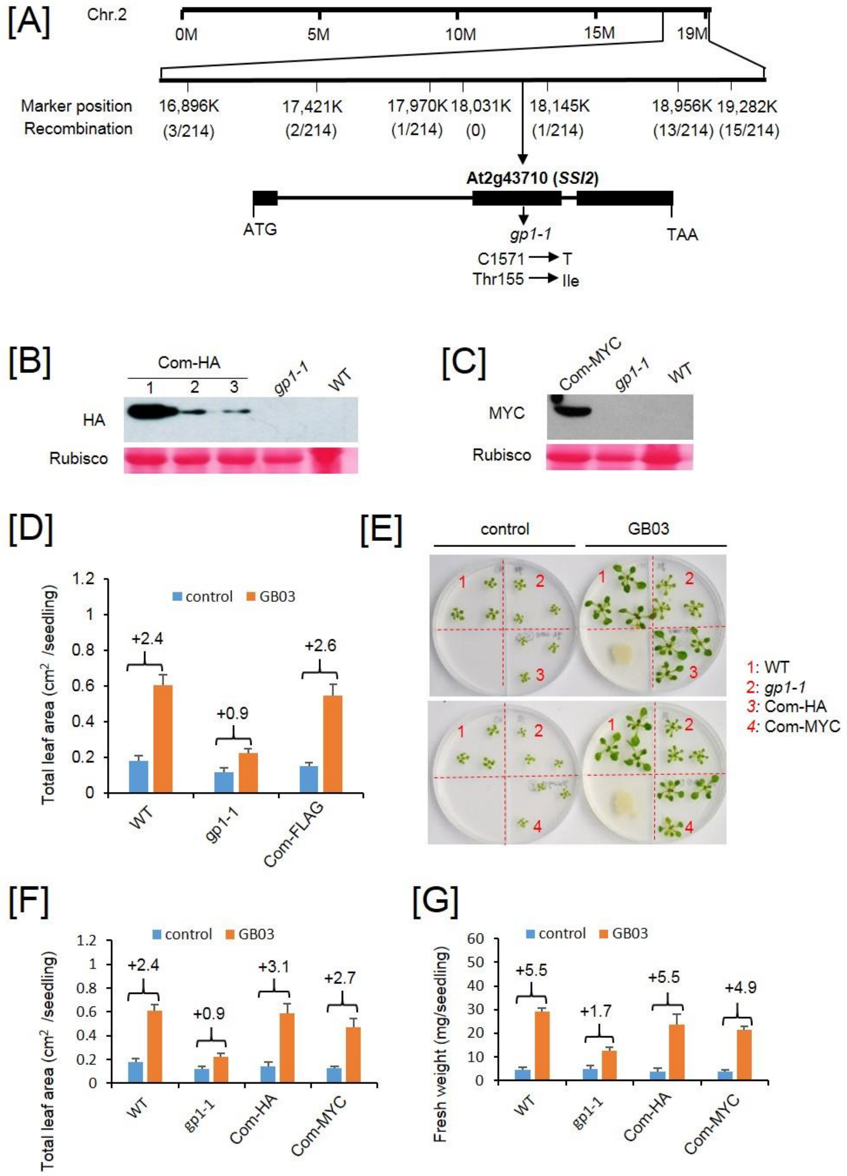


**Figure S2. Transgenic expression of At2g43710 in *gp1-1* rescues plant inducible vigor. [A]** Map-based cloning of *gp1-1* identified At2g43710. The chromosome 2 coordinate, markers positions, the recombination frequencies and the mutation site of At2g43710 were shown. **[B]** Transgenic expression of $Pro_{GP1}$::GP1-HA in *gp1-1* (Com-HA) restored protein accumulation of GP1, as shown in the western blot. **[C]** Transgenic expression of $Pro_{GP1}$::GP1-MYC in *gp1-1* (Com-MYC) restored protein accumulation of GP1. **[D]** Total leaf area per seedling quantified at 9 DAT. Mean ± SD (n=6). Fold changes are shown above each group. **[E]** Transgenic expression of $Pro_{GP1}$::GP1-HA (Com-HA) or $Pro_{GP1}$::GP1-MYC (Com-MYC) in *gp1-1* restored plant inducible vigor. Images were taken at 9 DAT. Red dotted lines indicate plastic partitions. **[F] and [G]** Total leaf area per seedling **[F]** and whole-seedling fresh weight **[G]** were quantified at 9 DAT. Mean ± SD (n=6). Fold changes are shown above each group. Related to Figure 2.

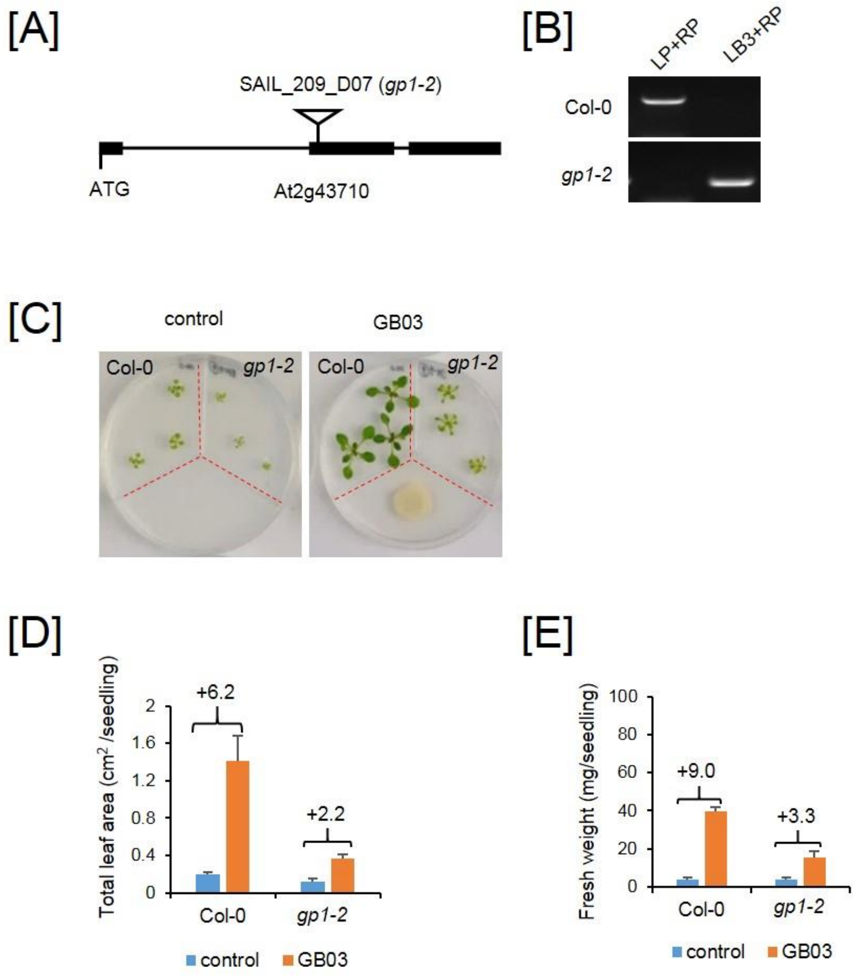


**Figure S3. The impairment of inducible vigor in the *gp1* mutants is caused by At2g43710.** **[A]** A schematic illustration of the T-DNA insertion site of *gp1-2* (SAIL_209_D07). **[B]** Gel image of the *gp1-2* genotyping result by PCR. **[C]** Comparison of *gp1-2* and Col-0 in response to GMVs. Plants were grown in 1/2-strength MS medium. Red dotted lines indicates plastic partitions in the petri dishes. Images were taken at 9 DAT. Quantification of total leave area per seedling and whole-seedling fresh weight are shown in **[D]** and **[E]**, respectively, Mean ± SD (n = 9). Fold changes are shown above each group. Related to Figure 2.

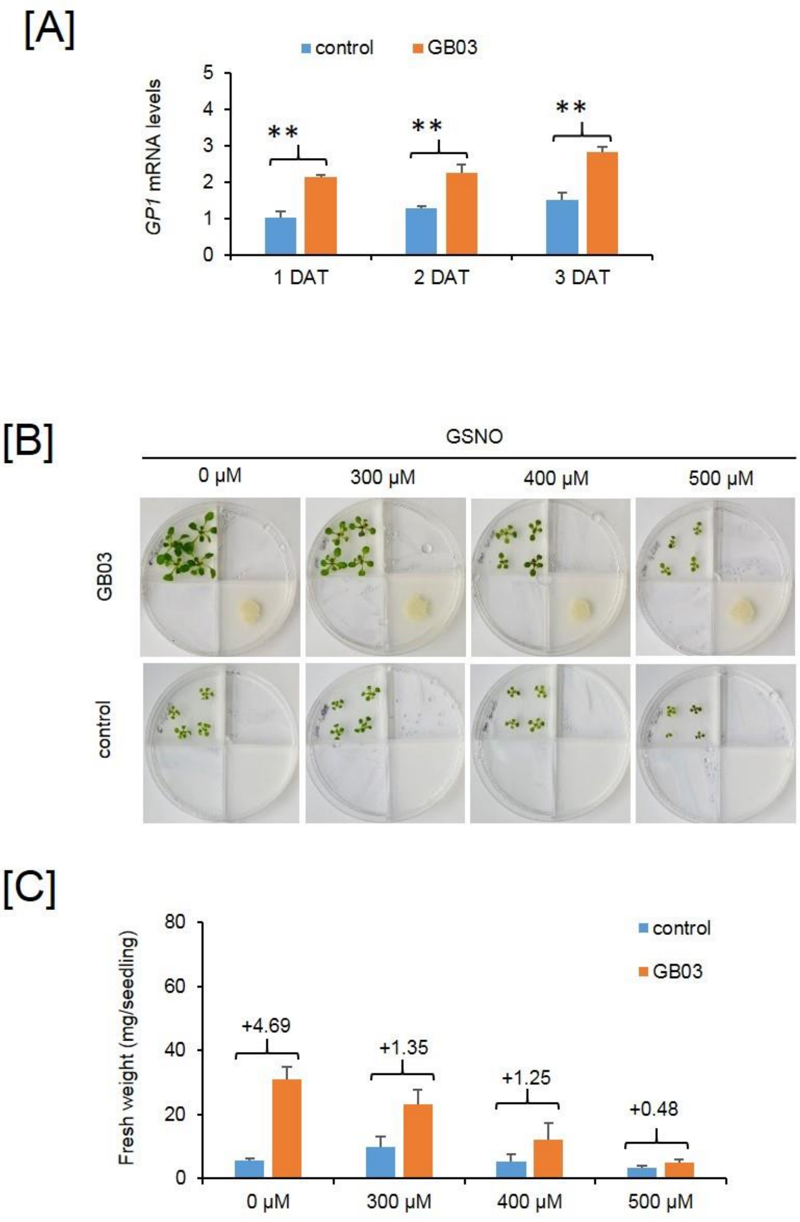


**Figure S4. Exogenous application of nitric oxide (NO) impairs plant inducible vigor. [A]** Relative gene expression levels of *GP1* in the WT seedlings after GMV treatment for 1, 2, 3 days; Values are normalized to *GP1* expression level in the control plants at 1 DAT. Mean ± SE (n=3). ** $p < 0.01$, student t-test; **[B]** Phenotype of WT seedlings grown in 1/2-strength MS containing 0 µM, 300 µM, 400 µM, and 500 µM GSNO, which is a NO donor. Images were taken at 10 DAT. Quantification of whole-seedling fresh weight is shown in **[C]**, mean ± SD (n = 8). Fold changes are shown above each group. An asterisk indicates $p < 0.05$, Student t-test. Related to Figure 2.

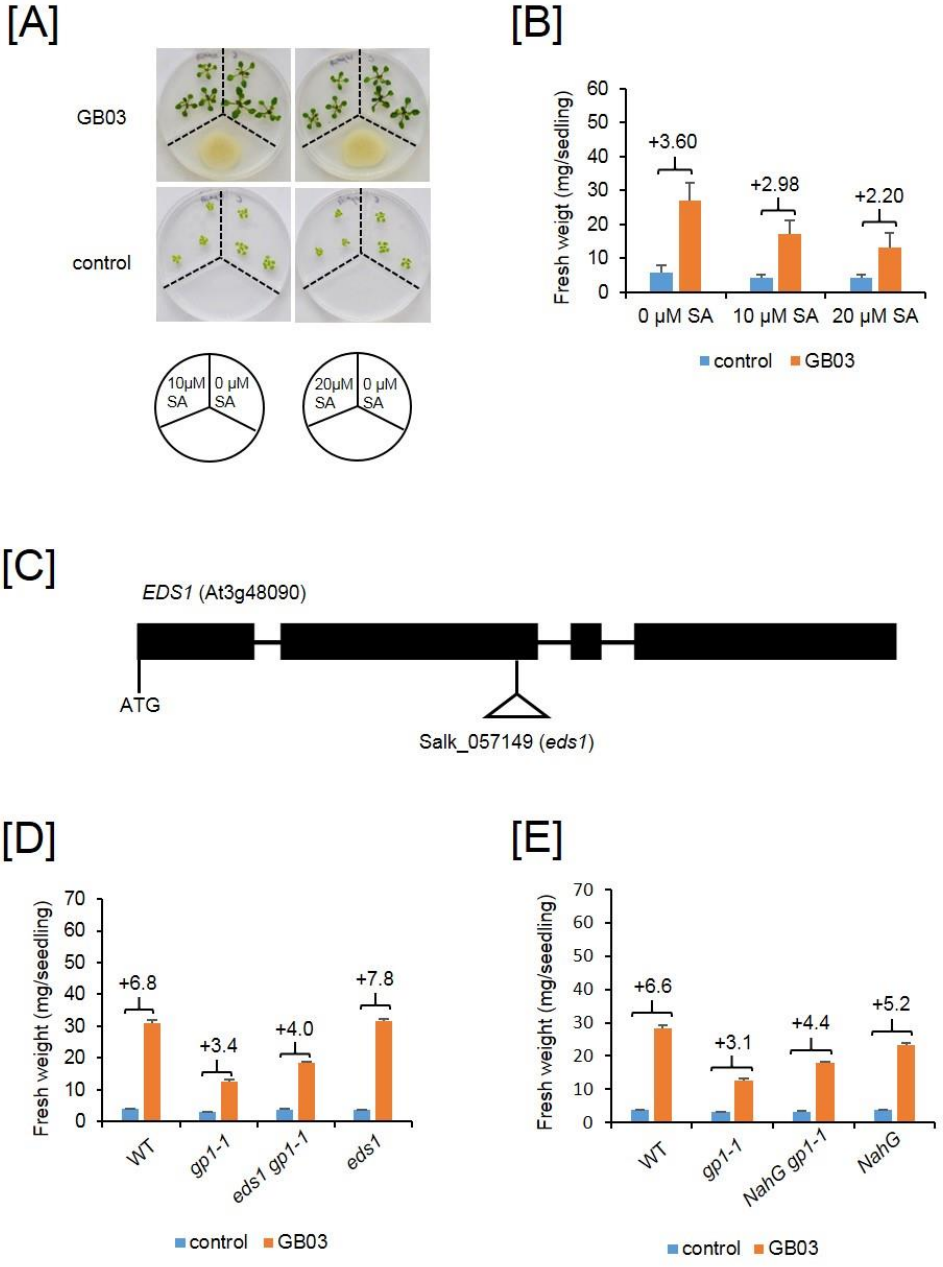


**Figure S5. The reduction of plant inducible vigor in *gp1* mutants partially depends on SA-mediated defense. [A]** Phenotype of WT seedlings grown in 1/2-strength MS containing 0 μM, 10 μM and 20 μM SA. Images were taken at 11 DAT. Quantification of whole-seedling fresh weight is shown in **[B]**. Mean ± SD (n = 9). Fold changes are shown above each group. **[C]** A schematic illustration of the T-DNA insertion site of the *eds1* mutation (SALK_057149). **[D]** Fresh weight per seedling was quantified for WT, *gp1-1*, *eds1 gp1-1*, and *eds1* at 10 DAT. Mean ± SD (n=8). **[E]** Fresh weight per seedling was quantified for WT, *gp1-1*, *NahG gp1-1*, and *NahG* at 10 DAT. Mean ± SD (n=8). Fold changes are shown above each group. Related to Figure 3.

[A]

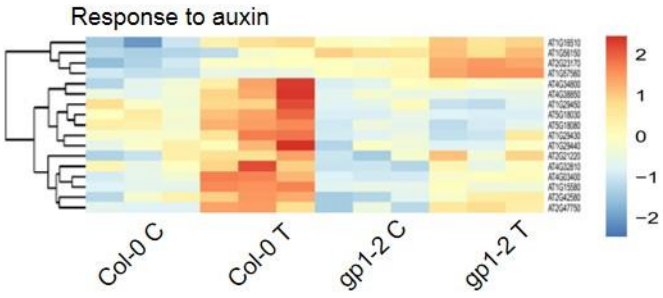


[B]

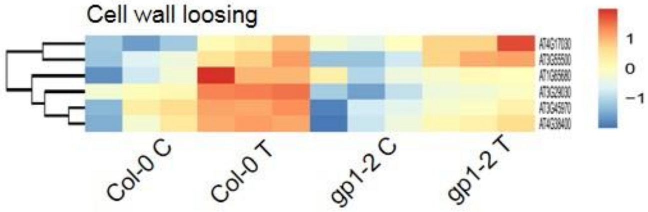


[C]

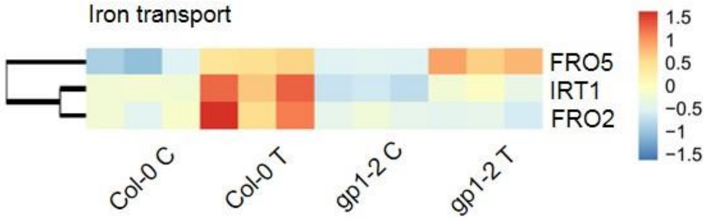


**Figure S6. GMVs induce the expression of growth-related genes in Arabidopsis (Col-0).** Heatmaps of GMV-induced DEGs that are involved in auxin response **[A]**, cell wall loosing **[B]** and Iron transport **[C]** are shown. Detailed gene lists were shown in Dataset S1. Related to Figure 4.

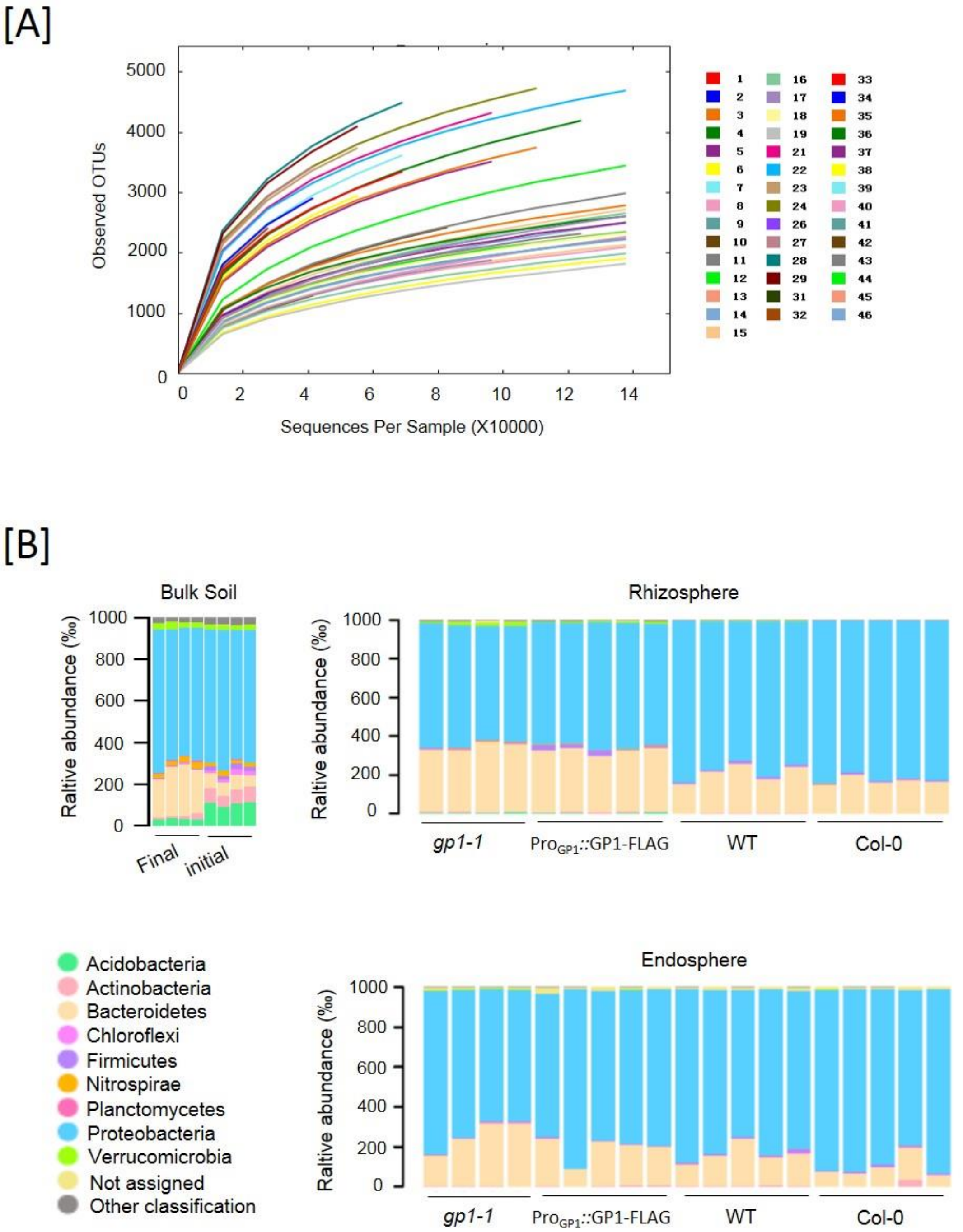


**Figure S7. GP1 dysfunction alters the assemblage of root-associated rhizobacteria community. [A]** Rarefaction analysis. Rarefaction curves are based on all the quality sequences obtained with a depth of 22843-690623 sequences per sample. The OTUs were defined on co-clustered quality sequences and rarefaction analysis was performed for all the samples. The color keys indicate different samples (see Dataset S2 for the detailed sample names). **[B]** Taxonomic structure of the ACM at phylum level in bulk soil, rhizosphere and root samples. Related to Figure 5.

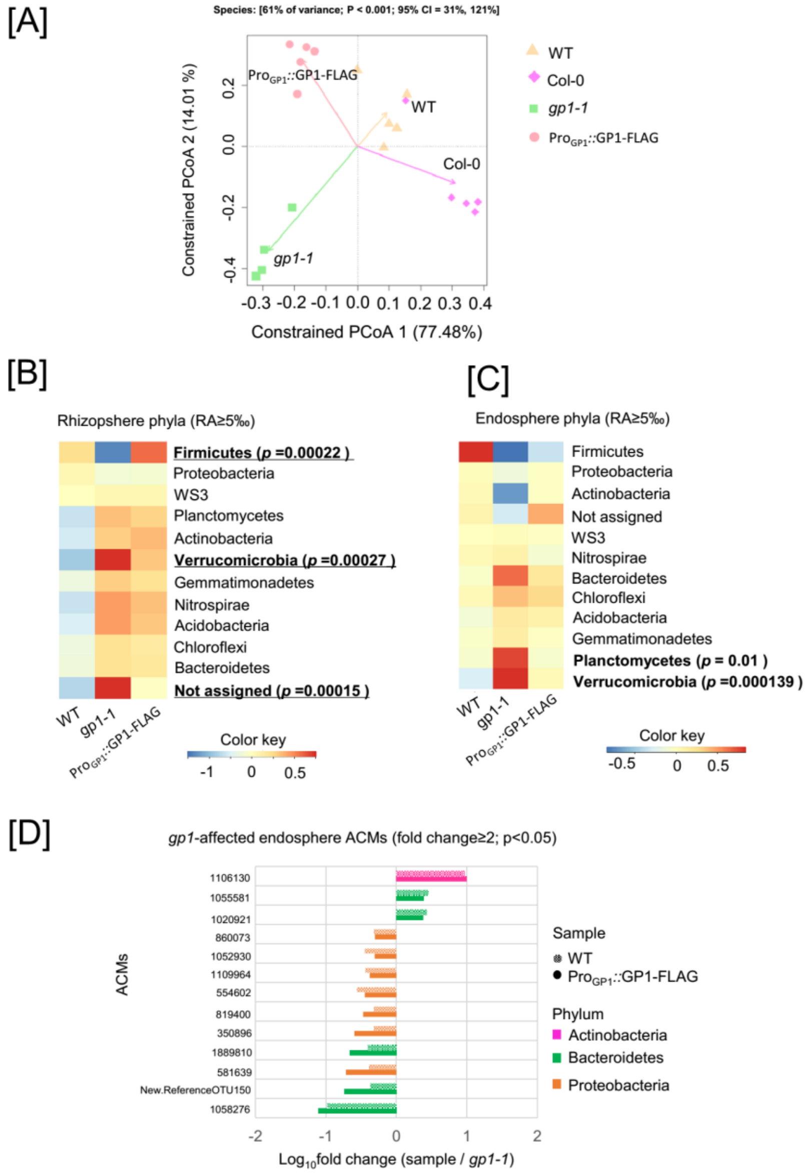


**Figure S8. GP1 dysfunction alters the assemblage of root-associated rhizobacteria community. [A]** Constrained principal coordinate analysis (PCoA) of ACM in the rhizosphere of WT, *gp1-1*, Col-0 and Pro$_{GP1}$*::*GP1-FLAG. The percentage of variation explained by each axis refers to the fraction of the total variance of the data (ACM) explained by the constrained factor. **[B]** Heatmap showing the relative abundances of rhizosphere phylums (RA≥5‰) in WT, *gp1*-1 and Pro$_{GP1}$*::*GP1-FLAG/*gp1-1*. The bold indicates *gp1*-affected phylums (WT VS *gp1-1* P<0.05 and Pro$_{GP1}$*::*GP1-FLAG/*gp1-1* VS *gp1-1* P<0.05); **[C]** Heatmap showing the relative abundances of endosphere phylums (RA≥5‰) in WT, *gp1-1* and Pro$_{GP1}$*::*GP1-FLAG/*gp1-1*. The bold indicates *gp1*-affected phylums (WT VS *gp1-1* P<0.05 and Pro$_{GP1}$*::*GP1-FLAG/*gp1-1* VS *gp1-1* P<0.05) **[D]** ACMs that are significantly changed in *gp1-1* mutant and complemented by Pro$_{GP1}$*::*GP1-FLAG in the endosphere compartment, as determined by fold changes of ACM abundance in WT and Pro$_{GP1}$*::*GP1-FLAG versus *gp1-1* (RA≥5‰; Fold change ≥ 2, Tukey p<0.05). Detailed RA and statistical analysis are listed in Dataset S2. Related to Figure 5.

**Table S1.**

Primers used in this study

| primer name | sequence | aims | from |
|---|---|---|---|
| SAIL_209_D07-LP | AACAGACATTGAGGAAAGCTTC | for *gp1-2* genotyping | |
| SAIL_209_D07-RP | CTTTTCGATCTGCCTCATGTC | | |
| SK069657-LP | CGGCTGTCATTTCTCTATTGC | for *act1-5* genotyping | |
| SK069657-RP | CTTGGCTGTATGCTTCTTTCG | | |
| SK057149-LP | TCACAGGACATTCCTCAGGAG | for *eds1* genotyping | |
| SK057149-RP | TTTATGGGCTTGACACTTTGG | | |
| NahG-F | GCCTTAGCACTGGAACTCTG | for NahG genotyping | |
| NahG-R | TCGGTGAACAGCACTTGCAC | | |
| LBb1.3 | ATTTTGCCGATTTCGGAAC | for salk lines genotyping | |
| LB3 | TAGCATCTGAATTTCATAACCAATCTCGATACAC | for sail lines genotyping | |
| At2g43710-EcoR1-F | GGGAATTCGGTCGTCCTAGCTGAATGGAC | for construct of the complementation test | |
| At2g43710-Xba1-R | GCTCTAGAGAGCTGCACTTCTCTGTCGTG | | |
| SSI2-qF | CTTTCAGATCTCCCAAGTTCCT | for Q-PCR of At2g43710 | |
| SSI2-qR | GGTGGCGTAAATGGTTTCTTC | | |
| PR1-qF | GGTTAGCGAGAAGGCTAACTAC | for Q-PCR of PR1 | |
| PR1-qR | CATCCGAGTCTCACTGACTTTC | | |
| ACT2-qF2 | TGTGTGACAAACTCTCTGGG | for Q-PCR of Actin2 | |
| ACT2-qR2 | GGCATCAATTCGATCACTCAG | | |
| 799F-B | ggagtgagtacggtgtgcAACMGGATTAGATACCCKG | 16S rRNA site specific PCR primers with bridging sequence | *(45, 46)* |
| 1193R-B | gagttggatgctggatggACGTCATCCCCACCTTCC | 16S rRNA site specific PCR primers with bridging sequence | *(45, 46)* |
| FP-1 | ACTCTTTCCCTACACGACGCTCTTCCGATCTgcttGCGTtggagtgagtacggtgtgc | Barcode primers (barcode sequence marked in red) | *(47)* |
| FP-2 | ACTCTTTCCCTACACGACGCTCTTCCGATCTgcttGTAGtggagtgagtacggtgtgc | Barcode primers (barcode sequence marked in red) | *(47)* |
| FP-3 | ACTCTTTCCCTACACGACGCTCTTCCGATCTgcttACGCtggagtgagtacggtgtgc | Barcode primers (barcode sequence marked in red) | *(47)* |
| FP-4 | ACTCTTTCCCTACACGACGCTCTTCCGATCTgcttCTCGtggagtgagtacggtgtgc | Barcode primers (barcode sequence marked in red) | *(47)* |
| FP-5 | ACTCTTTCCCTACACGACGCTCTTCCGATCTgcttGCTCtggagtgagtacggtgtgc | Barcode primers (barcode sequence marked in red) | *(47)* |
| FP-6 | ACTCTTTCCCTACACGACGCTCTTCCGATCTgcttAGTCtggagtgagtacggtgtgc | Barcode primers (barcode sequence marked in red) | *(47)* |
| FP-7 | ACTCTTTCCCTACACGACGCTCTTCCGATCTgcttCGACtggagtgagtacggtgtgc | Barcode primers (barcode sequence marked in red) | *(47)* |
| RP-A | GACTGGAGTTCAGACGTGTGCTCTTCCGATCTctgtGCGTtgagttggatgctggatgg | Barcode primers (barcode sequence marked in red) | *(47)* |
| RP-B | GACTGGAGTTCAGACGTGTGCTCTTCCGATCTctgtGTAGtgagttggatgctggatgg | Barcode primers (barcode sequence marked in red) | *(47)* |
| RP-C | GACTGGAGTTCAGACGTGTGCTCTTCCGATCTctgtACGCtgagttggatgctggatgg | Barcode primers (barcode sequence marked in red) | *(47)* |

| | | | |
|---|---|---|---|
| RP-D | GACTGGAGTTCAGACGTGTGCTCTTCCGATCTct gtCTCGtgagttggatgctggatgg | | |
| RP-E | GACTGGAGTTCAGACGTGTGCTCTTCCGATCTct gtGCTCtgagttggatgctggatgg | | |
| RP-F | GACTGGAGTTCAGACGTGTGCTCTTCCGATCTct gtAGTCtgagttggatgctggatgg | | |
| RP-G | GACTGGAGTTCAGACGTGTGCTCTTCCGATCTct gtCGACtgagttggatgctggatgg | | |
| RP-H | GACTGGAGTTCAGACGTGTGCTCTTCCGATCTct gtGATGtgagttggatgctggatgg | | |
| 2P-F | AATGATACGGCGACCACCGAGATCTACACTCTTT CCCTACACGACGCTCTT | Index primers (index sequence marked in blue) | *(47)* |
| 2P-R | CAAGCAGAAGACGGCATACGAGATAGTGACCTGA CTGGAGTTCAGACGTGTGCTCTT | | |

**Data S1. (separate file)**

DEG (differentially expressed gene) lists generated from the RNA seq data. This dataset contains multiple sheets. The titles of individual DEG lists are summarized in the first sheet named "Contents".

**Data S2. (separate file)**

Micobiota datasets. This dataset contains the sample number, genotypes, barcode sequences, OTU tables of TIC, ACM, and relative abundance at phylum, family, and ACM levels. A summary of the individual sheets is listed in the first sheet named "Contents".
